# Strong effects of soil microbes on early-life defense alter phenotypic correlations in plant leaves

**DOI:** 10.64898/2026.06.29.735305

**Authors:** Eunnuri Yi, Mikhaela Ferguson, Maria Altamirano Chavez, Corlett W. Wood

**Author notes:** Both authors contributed equally (Eunnuri Yi, Mikhaela Ferguson).

## Abstract

Microbes have wide-ranging effects on host phenotype. However, whether these effects extend to the relationships among host traits remains unknown. We tested whether microbes affect phenotypic correlations among early life traits in the weedy legume *Medicago lupulina.* In a field common garden, we inoculated plants with two types of microbes: nitrogen-fixing bacteria and rhizosphere microbial communities. We found that microbes modify phenotypic integration (phenotypic correlations) in leaves, primarily due to effects on independent effects on leaflet area and trichome density, a defense trait. While microbial effects on leaflet size were associated with overall plant growth, variation in trichome density was decoupled from growth and not predicted by investment in mutualism. Our results highlight the unique insights that can come from a multivariate approach to organismal plasticity, and raise the intriguing possibility that microbes may have an outsized impact on defense traits and their integration with organismal function.

## Introduction

Many plant and animal traits are jointly determined by microbes ((Friesen et al. 2011; O’Brien et al. 2019, 2021)). A diverse array of microbes affect host phenotype, from coevolved endosymbionts—like *Wolbachia* and *Buchnera* in insects and *Vibrio* in squid—to entire microbial communities (Heath-Heckman et al. 2014; Hawkes et al. 2021; O’Brien et al. 2021; Higashi et al. 2024; Douglas 1998). However, organisms are not simply collections of individual traits; they are composed of integrated suites of traits that work together to perform specific functions. The observation that microbes have far-reaching effects on their hosts raises an important question: do microbial effects extend beyond individual traits to impact phenotypic integration in their hosts?

The relationships among functionally related traits play a central role in adaptation and evolution (Lande and Arnold 1983; Pigliucci 2003; Arnold et al. 2008). Phenotypic correlations can arise from natural selection acting to integrate functionally related traits, or from allocation tradeoffs across competing functions (De Jong 1993; Conner and Sterling 1996; Murren 2002; Damián et al. 2020; McGlothlin et al. 2022). Correlations among traits influence evolutionary trajectories by generating correlated responses to selection (Schluter 1996; Chenoweth et al. 2010). However, the relationships among traits are not static. Trait correlations are often highly sensitive to the environment (reviewed in Sgrò and Hoffmann 2004; Wood and Brodie 2015). Correlations among traits frequently strengthen, weaken, or even reverse across environments. These effects arise from a wide range of abiotic and biotic factors, including temperature, moisture, light, predation, competition, diet, and resource availability (reviewed in Wood and Brodie 2015).

Microbial symbionts are conspicuously absent from this literature, despite their profound effects on host phenotype and growing evidence that the microbial environment is as heterogeneous as the physical environment. There is substantial spatial variation in the composition and function of entire microbial communities (Wagner et al. 2014; Petipas et al. 2020; Lange et al. 2023). And even in tightly coevolved symbioses, symbiont genotypes differ substantially in their effects on host traits and fitness (Sachs et al. 2009; Heath and Stinchcombe 2014; Riley et al. 2023; Gano-Cohen et al. 2019). However, microbial effects on trait integration cannot be inferred from the extensive literature on microbe-mediated plasticity, even those studies that report microbial effects on multiple host traits (Martin et al. 2022). This is because changes in trait means do not necessarily alter trait correlations. For microbes to alter trait correlations, individual hosts must differ in their responses to microbes. As a result, we know surprisingly little about whether pervasive microbial effects on individual host traits extended to the *relationships* among traits.

Nor can we extrapolate from the broader literature on environmental effects on trait integration, because microbes simultaneously alter multiple aspects of host biology. Microbes provide and consume resources, induce local and systemic immune responses, modify stress tolerance, and accelerate or delay host ontogeny (Jung et al. 2012; Pringle 2016; O’Brien et al. 2021; Fontaine et al. 2022; Bogar 2024). A multivariate perspective on microbe-mediated plasticity is therefore essential to identify traits that are unusually sensitive to microbes; determine whether microbes induce predictable or novel combinations of host traits; and to evaluate whether microbes strengthen, relax, or modify phenotypic integration. The few studies that have tackled microbe-mediated plasticity from a multivariate perspective demonstrate that this happens, and the new insights that arise from this approach (Wood et al. 2026; Zhao et al. 2026).

Here, we investigated how microbes affect relationships among early-life traits in black medick (*Medicago lupulina*), a widespread roadside weed. We focused on two well-characterized axes of phenotypic variation in plants: leaf and growth traits. Leaves, the primary site of photosynthesis, are a single organ made up of several specialized tissues. We measured five leaf traits that are associated with the leaf economic spectrum, which organizes multivariate leaf phenotypes along a spectrum from resource acquisitive to resource conservative (Donovan et al. 2011a; Reich 2014; Díaz et al. 2016; Anderegg et al. 2018). Growth traits, such as biomass and measurements of length or height, are a common bioassay for microbial effects on hosts (Vessey 2003; Friesen et al. 2011; O’Brien et al. 2021; Rahman et al. 2023). Growth traits tend to be strongly positively correlated: larger individuals are larger in all dimensions (Díaz et al. 2016).

We inoculated plants with two different kinds of soil microbes: nitrogen-fixing bacteria (rhizobia) and an intact soil microbial community sampled from the rhizosphere of wild plants. Rhizobia are a textbook resource-exchange mutualism that fix atmospheric nitrogen in plant root nodules (Mylona et al. 1995; Masson-Boivin and Sachs 2018). The rhizosphere microbial community comprises diverse microbial taxa that live in association with plant roots. In our experiment, its exact composition was unknown, but it likely included species whose effects on the host plant range from mutualistic to commensal to parasitic (Prashar et al. 2014; Ling et al. 2022). We compared phenotypic correlations in plants inoculated with rhizobia or rhizosphere microbes to a microbe-depleted control, a common approach in experiments testing microbe-mediated plasticity.

We asked two questions: (1) Do microbial treatments alter phenotypic correlations among leaf and growth traits?; and (2) Do microbes induce predictable or unique trait combinations in their hosts? We found that microbes significantly strengthened phenotypic correlations in leaves relative to the microbe-depleted control, and altered the direction of phenotypic correlations. Unique combinations of leaf traits distinguished different microbial treatments. By contrast, microbes had a comparatively weak effect on phenotypic correlations in growth traits. We identified the two leaf traits that drove microbial effects on phenotypic integration in leaves—leaflet size and the density of defensive hairs (trichomes)—and tested whether microbial effects on these traits were explained by growth or investment in the rhizobia mutualism. While microbial effects on leaflet size were associated with overall plant growth, variation in trichome density was decoupled from growth and not predicted by investment in mutualism. Our results highlight the unique insights that can come from a multivariate approach to organismal plasticity. Finally, our results raise the intriguing possibility that microbes may have an outsized impact on defense traits, and their integration with other organismal functions.

## Methods

### Study system

Black medick (*Medicago lupulina*) is a common weed naturalized to North America abundant throughout temperate and subtropical regions throughout the world, thriving in disturbed areas such as roadsides and lawns (Turkington and Cavers 1979). It is primarily self-fertilizing, although it is also capable of outcrossing. It is a short-lived perennial, but often flowers in its first year, approximately 6-8 weeks after germination in benign conditions (Turkington and Cavers 1979, Wood personal observation). *Medicago lupulina* primarily partners with rhizobia in the genus *Ensifer* (or *Sinorhizobium)*, especially *E. meliloti* and *E. medicae* (Harrison et al. 2017a, 2017b).

### Field experiment

We compared microbe-depleted and microbe-inoculated plants by germinating seeds and planting them in five different treatment groups. These groups consisted of one microbe-depleted group, two groups inoculated with pure cultures of different rhizobia strains, and two groups inoculated with intact soil communities. Each group contained between 20-25 plants, for a total of 121 plants. Plants were grown in the deer exclusion plot beside the greenhouse at Mountain Lake Biological Station (MLBS), Salt Pond Mountain, Giles County, Virginia, USA, 37.3746 N, −80.5214 W, at an elevation of 1,160 meters.

To germinate the *Medicago lupulina* seeds, the seed coat was gently removed using sandpaper. The seeds were then scarified using a razor blade under a dissecting microscope. Scarified seeds were sterilized by placing them in ethanol for 30 seconds, then in bleach for one minute, before rinsing them with water and soaking in water for 30 minutes. Scarified, sterilized seeds were then placed into the respective soil treatment containers.

All soil was autoclaved to kill the microbes present and ensure property changes within the soil were consistent amongst groups. Rhizobia nodules were taken from roots of *Medicago lupulina* found at MLBS and at Pandapas Pond, a lower elevation site. Nodules were crushed and grown on solid culture media. After 2-3 days, the bacteria were transferred to liquid media. 1mL of liquid media was dispensed into the soil to inoculate each container of rhizobia treatment soil containers. To inoculate the community plants, soil samples from different sites were into the autoclaved soil. The community soil sample comprised 10-20% of the total soil volume in the container. Germinated, sterilized seeds from the same plant were planted in each container. All of the seeds were collected from the same plant to ensure genetic similarity.

Our experimental approach was successful at minimizing rhizobia contamination and manipulating microbes in the field. Very few plants in the microbe-depleted treatment formed nodules, indicating that rhizobia contamination was minimal (Figure 4A, Table S3B).

### Trait measurements

All plants studied were assessed on the following traits: growth rate and above- and belowground biomass; leaflet mass, area, and leaflet mass area; simple trichome density; stomata density; and chlorophyll content. We measured the length of the longest stem every 2-3 days throughout the experiment as a proxy for growth rate. Plants were harvested approximately three weeks after planting, after which we weighed dried shoots and roots to measure above- and belowground biomass. Leaflet mass area and growth rate were used to assess investment in creation of photosynthetic structures. We sampled the center leaflet of the topmost fully expanded trifoliate leaf on the longest/central stem. Leaflet mass area was calculated by scanning on a leaflet per plant in an office scanner with a ruler to calibrate and using *ImageJ* software to calculate area via number of pixels (Schneider et al. 2012). Those leaflets were collected and weighed for dry mass. The growth rate of one leaflet per plant was measured throughout the experiment by measuring the length and width with a ruler. The widest part of the leaf was used to standardize the width measurement.

Simple trichome density was obtained by placing the underside of the leaf under a dissecting microscope. A ruled microscope slide with areas of 1mm^2^ marked was placed over the leaf. The grid block to the right of the stem at the widest section of the leaf was counted for simple trichome density per mm^2^.

Chlorophyll content represented how much the plant was investing in performing photosynthesis. Stomata density represents investment in photosynthetic structures and potential to perform gas exchange to provide resources for photosynthesis. To collect data on stomata density, a mold of the underside of the leaf was made using *NewSkin* liquid bandage (Advantice Health, 2023). Clear tape was used to transfer the mold to a microscope slide. The leaf mold was put under a compound microscope. A picture was taken of the mold under the microscope and the image was put into *ImageJ* to count the number of stomata present (Schneider et al. 2012). Relative chlorophyll abundance was obtained using a *SPAD 502 Chlorophyll Meter* (Konica Minolta, Inc., 2009). Leaves were placed in the cuvette of the machine, which used transmittance of light to measure how much chlorophyll was present.

### Statistical analysis: Microbial effects on plant traits and phenotypic integration

We performed all analyses in R version 4.6.0 and R Studio Version 2026.05.1+225 (R Core Team 2026). Unless otherwise specified, we fit all linear models using the package *glmmTMB* (Brooks et al. 2017) and tested significance using Wald chi-square tests and type III sums of squares in the *car* package, assuming a Gaussian distribution (Fox and Weisberg 2019). We verified that residuals were normally distributed and homoscedastic using the *DHARMa* package (Hartig 2026). We conducted post-hoc comparisons between treatment groups using Tukey HSD tests implemented in the package *emmeans* to estimate marginal means and confidence intervals (Lenth 2026), and created figures using the *ggplot2* package (Wickham 2016).

To test for microbial treatment effects on plant leaf traits (leaflet area, leaf mass area, chlorophyll, stomatal density, and trichome density), growth traits (shoot biomass, root biomass, shoot-to-root ratio), and nodule number, we compared these traits across microbial treatments. These models included microbial treatment as a fixed effect and rack number as a random effect to account for spatial variation. To address violations of model assumptions, we fit generalized linear mixed models with the appropriate error distribution (log-transformed for leaf area, and Gaussian for all other traits) (Table S3).

We performed separate analyses for a suite of leaf traits (leaf area, leaf mass per area, chlorophyll content, stomatal density, and trichome density) and a suite of growth traits (shoot biomass, root biomass, shoot:root ratio, and stem length). We also reran analyses on leaf traits excluding trichome density (leaf area, leaf mass per area, chlorophyll content, and stomatal density) to assess whether trichome density was the primary driver for the differences observed in leaf phenotypic correlations. Observations with missing values for the traits under analysis were excluded on a per-analysis basis.

### Statistical analysis: Multivariate microbial effects on trait covariances

To test whether soil microbes influenced total phenotypic covariation in plant leaf and growth traits, we compared phenotypic variance-covariance (**P**) matrices between microbial treatments and visualized their first and second principal components using the prcomp function from the core *stats* package (R Core Team, 2026). We performed these analyses on a suite of five leaf traits, a suite of four growth traits, and a modified suite of four leaf traits excluding trichome density. We compared covariance structure to the microbe-depleted control—as is common in the microbiome literature—to reduce multiple testing (Dunnett 1955; Day and Quinn 1989; Vorholt et al. 2017; Midway et al. 2020).

We mean-standardized all traits within treatments prior to analysis by dividing each individual’s raw value by its treatment-specific trait mean (Delahaie et al. 2017). We did so because trait standardization can have a major impact on a matrix comparison analysis, especially if the original traits were measured on different scales, as ours were (Hansen et al. 2011). Mean standardization removes differences in trait variation attributable to measurement scale, preserving the relative differences between different traits (Hansen et al. 2011; Houle et al. 2011; Delahaie et al. 2017). Variance standardization (i.e., dividing by the standard deviation) converts all traits into standard deviation units and thereby eliminates any differences in trait variation among treatments: thus it is not a viable option when comparing total phenotype variation, as we do below. We also produced correlation matrices after variance-standardizing and mean-centering traits within each treatment to look directly at trait correlations (Table S12).

To test whether total phenotypic variation differed between microbial treatments, we compared the volume of **P** across treatments (Wood and Brodie 2015). We calculated the volume of **P** for each treatment by summing the eigenvalues for each matrix. We calculated the proportional difference in volume for each pair of P-matrices by calculating the difference in volume and dividing by the average volume of the two matrices.

To test whether the magnitude of trait correlations differed between microbial treatments, we compared the proportion of phenotypic variation captured by the major axis of phenotypic variation (**p_max_**; the first principal component). We calculated the proportion of variation on **p_max_** by dividing the dominant eigenvalue by the sum of all eigenvalues: 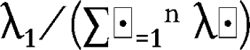.

To test whether the orientation of trait correlations differed between microbial treatments, we calculated the angle between **p_max_** for each pair of matrices. We extracted the dominant eigenvector (i.e., the first principal component) from each matrix, and calculated the angle between the dominant eigenvectors for each pair of matrices as in (Wood and Brodie 2015):

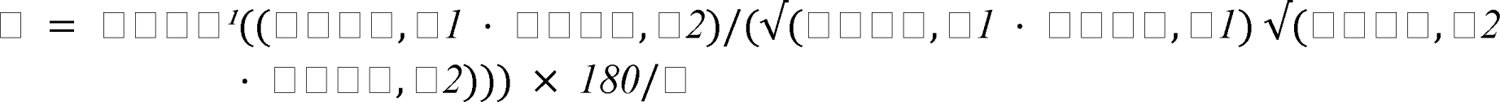

An angle of 0° means that there is no difference in the orientation of the major axis of phenotypic variation between two matrices, while an angle of 90° means that the major axes of phenotypic variation in the two matrices are orthogonal to each other. We took the absolute value of the cosine before calculating the inverse cosine, such that vectors differing by 180°—oriented in the same direction with different signs—became identical and all angles were between 0° and 90°.

We used permutation tests to determine which pairwise differences in total phenotypic variation, the magnitude of trait correlations, or the orientation of trait correlations were significant. To generate null distributions, we randomly sampled from our original unstandardized data without replacement to create two new groups. The sample size of each new group was determined by randomly drawing the sample size of one of the five microbial treatments. We then mean-standardized traits and calculated P-matrices for each new group and calculated the three matrix comparison metrics for each pair. We repeated this process 10,000 times. For differences in total phenotypic variation and the proportion of variation on **p_max_**, we considered empirical values that fell in the lower and upper 2.5% tails of these null distributions statistically significant (i.e., a two-tailed test). For the angle between **p_max_**, we considered empirical values that fell in the upper 5% tail of the null distributions statistically significant (i.e., a one-tailed test). We performed these permutation tests separately on leaf and growth trait matrices to generate separate null distributions for these analyses.

In addition to our tests of microbial effects on phenotypic variance and covariance for each treatment group compared to the microbe-depleted control, we also tested for differences in the multivariate mean phenotype between all treatment groups using multivariate analysis of variance (MANOVA). We ran three separate analyses: one that included the five leaf traits listed above, one that included the four leaf traits excluding trichome density, and one that included the four growth traits. We used Pillai’s trace to test significance because it is robust to violations of MANOVA assumptions such as multivariate normality and homogeneity of variance (Olson 1976; Finch 2005). To visualize multivariate differentiation among treatments and identify the traits which contributed most strongly towards the primary axes of visualization, we conducted linear discriminant analyses (LDA) on standardized trait values using the MASS package (Venables and Ripley 2002). We assessed the contribution of traits by extracting the unadjusted structure coefficients using the *candisc* package, which we visualized as plotted trait loadings on the first two linear discriminants (Friendly and Fox 2025) (Figure 3, Table 2).

### Statistical analysis: Tests of microbial effects via growth and nodulation

To test whether microbially-mediated effects on plant growth and overall performance corresponded to variation in focal leaf traits (leaflet area and trichome density), we used the prcomp function to extract the first principal component of the growth trait covariance matrix built across all treatments to generate a composite growth axis (R Core Team 2026, Figure S1, Table S1). We fit mixed-effects models for trichome density and leaf area as a function of the growth PC1, microbial treatment, and their interaction, with rack as a random effect. We compared the slopes of each treatment using the emtrends function (Lenth 2026).

To directly test whether the rhizobia mutualism affected focal leaf traits which exhibited the largest microbial effects, and whether this effect differed across microbial treatments, we fit models with leaf traits (leaflet area and trichome density) as the response variables and treatment, mean-centered nodule number, and their interaction as fixed effects. These models also included rack number as a random effect. We excluded the microbe-depleted treatment from this analysis.

Lastly, to assess whether the rhizobia mutualism overall facilitated growth and to test whether the per-nodule growth benefit derived from rhizobia differed across microbial treatments, we fit a model with log-transformed shoot biomass as the response and treatment, mean-centered nodule number, and their interaction as fixed effects. We included rack number as a random effect. A significant treatment-by-nodule number interaction would indicate that microbial communities impact the per-nodule benefit the plant derives from rhizobia. We excluded the microbe-depleted treatment from this analysis.

## Results

### Soil microbes modify leaf trait integration and correlations

Covariance matrix comparisons revealed that for leaf traits, microbes generally increased total phenotypic variation, strengthened phenotypic correlations, and sometimes modified the direction of phenotypic correlations; for growth traits, microbes did not overall significantly or consistently modify phenotypic correlations. Across all traits, we observed no consistent difference between rhizobia and community treatments in their effects on phenotypic integration across leaf and growth traits: both types modified trait relationships.

For leaf traits, both rhizobia treatments significantly increased the total variance (volume, or sum of eigenvalues) by 76.7-92.7%, while community treatments either had marginal or non-significant increases in volume (22.6-62.0%) compared to the microbe-depleted control (Figure 2F, Table S2). All four microbial treatments significantly strengthened phenotypic correlations: the proportion of variance represented by **p_max_** was 30.0-37.4% greater on microbial treatments compared to the microbe-depleted control (Figure 2G, Table S2). Thus microbial treatments affected the dimensional nature of the multivariate leaf phenotype. The proportion of leaf trait variance explained by **p_max_** was only 37% for the microbe-depleted control, whereas it ranged from 65.9-74.4% for all inoculated microbial treatments. The PC2 of inoculated microbial treatments represented just 12.4-17.5% of within-treatment variation, whereas the PC2 of the microbe-depleted control represented 24.6% of variation. One rhizobia and rhizosphere community treatment (both sampled from MLBS) also significantly modified the direction of phenotypic covariances (represented by the angle of **p_max_** compared to the microbe-depleted control), meaning that they reoriented the primary axis of leaf trait covariation by 36.9° and 63.8° respectively (Figure 2H; Table S2; Figure 2A-E).

**Figure 1:**
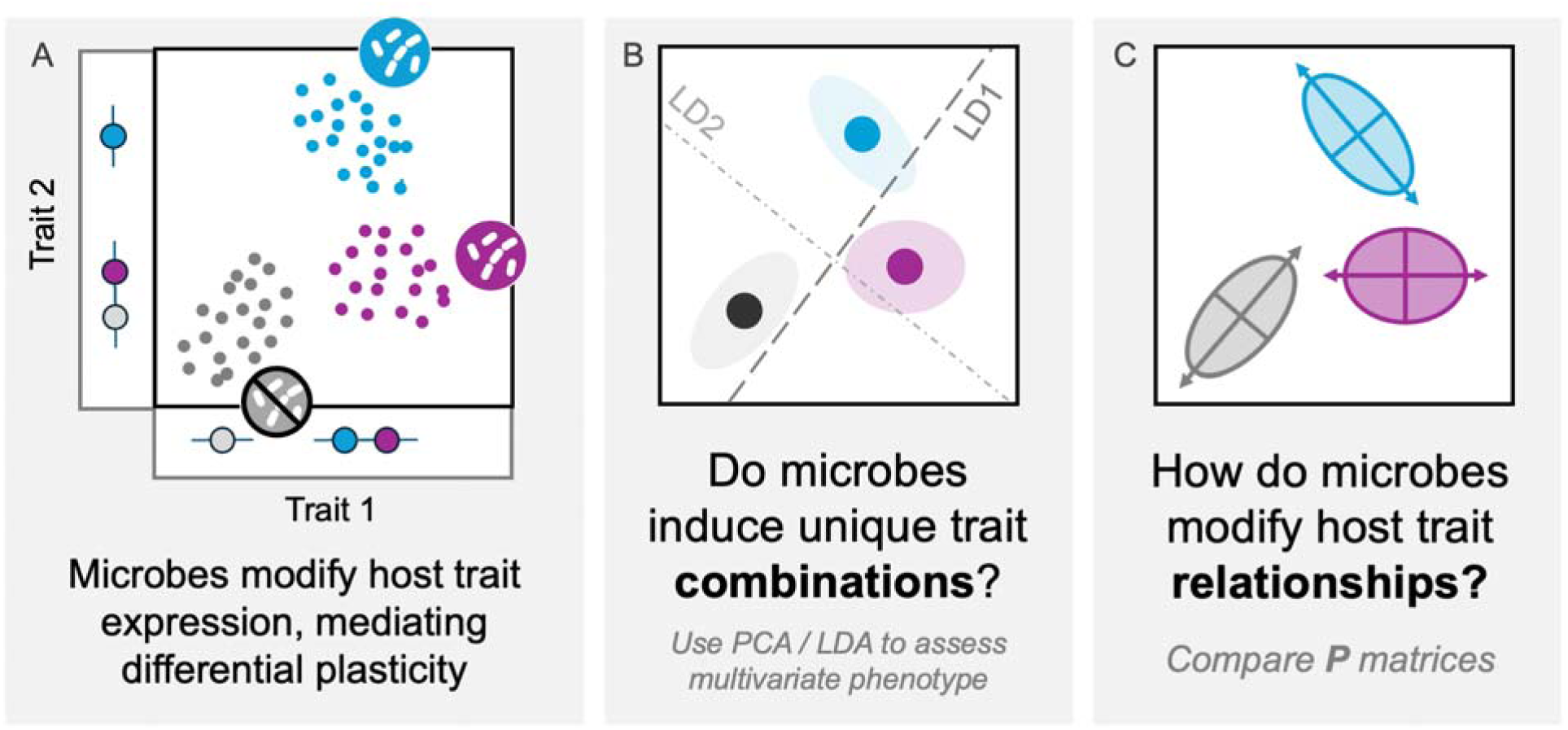
A multivariate perspective demonstrating potential microbial effects on multivariate host phenotypes and phenotypic integration. A) Microbial effects on host trait expression can be univariate or multivariate (modifying multiple traits simultaneously *and* differentially). B) If microbes induce unique trait combinations, they modify the multivariate phenotype; we represent mean effects on groups with centroids. Groups can be separated by LDA to identify trait combinations that separate treatments; PCA can identify overall axes driving variation. C) Microbes can also modify the trait *relationships* or correlations within hosts, which can be compared using P-matrices.

**Figure 2:**
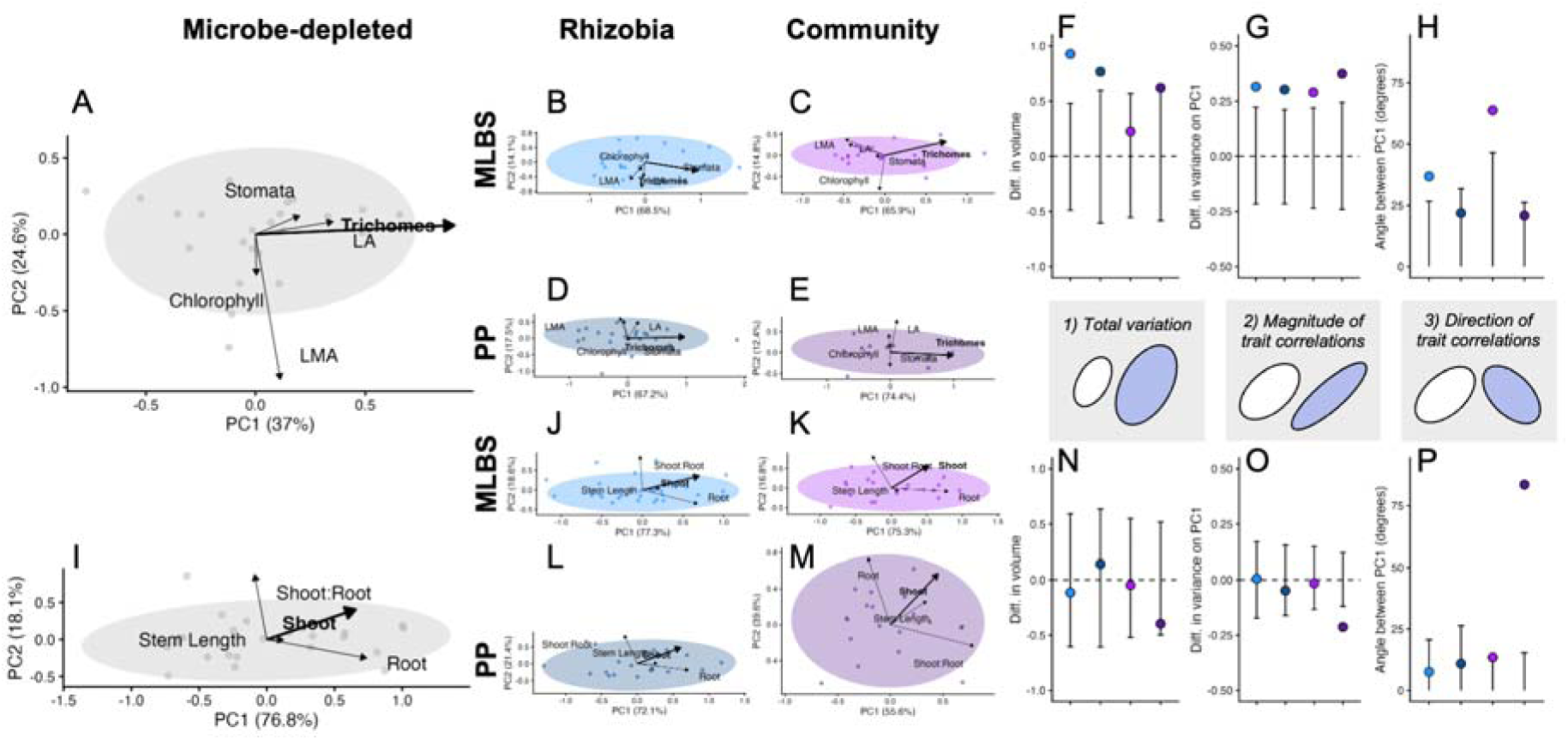
Principal Component Analyses and matrix comparisons of microbial treatments. Principal component analyses of leaf traits (A-E) and growth traits (F-J) across microbial treatments. Ellipses represent 95% confidence intervals. Panels K-P show matrix comparisons with permutation test 95% null distributions. Matrix comparison values are reported in Table S2.

Microbial treatments altered trait correlations. Different microbial treatments changed the strength and direction of trait correlations—most dramatically between trichome density and leaflet area (from −0.686 (community MLBS) to 0.375 (rhizobia PP))—but also between leaflet area and stomata density (from −0.209 (community PP) to 0.247 (microbe-depleted)), leaflet mass area and chlorophyll (−0.097 (community PP) to 0.758 (rhizobia PP)), leaflet mass area and stomata density (−0.375 (community PP) to 0.372 (rhizobia MLBS)), and leaflet mass area and leaflet area (−0.025 (microbe-depleted) to 0.580 (community MLBS), although these comparisons were not significance tested (Table S12). Most trait correlations changed across treatments to some degree, and not all changing trait correlations involved trichome density.

For leaf traits excluding trichome density, matrix comparisons showed that the strengthening of phenotypic correlations observed in the full analysis diminished: volume and proportion of variance effects largely fell within permutation boundaries. The community (PP) treatment actually exhibited a relative decrease in total volume compared to the microbe-depleted control (Table S9). Both community microbial treatments retained significant differences in the angle of overall trait correlations even after removing trichome density from the suite of leaf traits. MANOVA for the reduced leaf trait set remained significant (F □ □, □ □ □ = 5.407, P < 0.001; Table S8), and LDA for the four leaf traits (Figure S4I; Table S11) still produced comparable treatment discrimination with leaf area and chlorophyll as primary drivers (LD1: −0.981 and −0.442, respectively).

Microbes did not significantly or consistently modify phenotypic correlations in growth traits. Microbial effects on total phenotypic variation were not significant for growth traits, and only one community treatment (PP) had any significant effect on the proportion of variance on **p_max_**, which consistently represented a relatively large proportion of variance across all treatments (55-77%), reflecting a “size” axis explained primarily by shoot biomass (45-70%), root biomass (−21-80%), and moderately by stem length (10-34%) (Table 1). PC2 generally reflected allocation of growth represented by shoot:root ratio, with the exception of the community treatment from Pandapas Pond, which had a high shoot:root loading onto **p_max_** (0.7935; Table 1). Both community treatments modified the direction of phenotypic covariances, although the difference for the community (MLBS) treatment was not large (13.44° with a 95% permutation threshold of 12.57°; Table S2). The Pandapas Pond (PP) community treatment had relatively less variance represented on **p_max_**, meaning that it weakened phenotypic covariances. The community (PP) treatment also showed an unusual alignment of shoot:root ratio with its primary growth axis (Table 1), suggesting some disruption of growth integration—resulting in a dramatic change in the orientation of the axis of growth trait covariance—but this was unique to that single treatment. The trait loadings overall aligned in similar relative orientations for microbial treatments including the microbe-depleted control, with the exception of the Pandapas Pond community treatment.

**Table 1:** Trait loadings onto and variance explained by PC1 of each microbial treatment. Trait loadings represent the correlation between the original trait and PC1; the proportion of variance explained by PC1 for leaf and growth traits were calculated from covariance matrices with traits standardized by dividing by the group means.

|  | Microbe-depleted | Rhizobia (MLBS) | Rhizobia (PP) | Community (MLBS) | Community (PP) |
| --- | --- | --- | --- | --- | --- |
| **LEAF TRAITS** |  |  |  |  |  |
| *Proportion of Variation on PC1* | 0.3699 | 0.6852 | 0.6719 | 0.6594 | 0.7439 |
| Leaflet Area | 0.3540 | -0.0662 | 0.1890 | -0.4400 | 0.1078 |
| Leaflet Mass Area | 0.1134 | -0.2745 | -0.1389 | -0.4822 | -0.0253 |
| Chlorophyll | 0.0045 | -0.1122 | -0.0759 | -0.0691 | -0.0200 |
| Stomata Density | 0.2011 | -0.0372 | -0.0056 | -0.1149 | -0.0068 |
| Trichome Density | 0.9063 | 0.9520 | 0.9691 | 0.7456 | 0.9936 |
| **GROWTH TRAITS** |  |  |  |  |  |
| *Proportion of Variation on PC1* | 0.7684 | 0.7732 | 0.7205 | 0.7530 | 0.5556 |
| Shoot biomass | 0.6561 | 0.7000 | 0.6079 | 0.5222 | 0.4538 |
| Root biomass | 0.7387 | 0.6799 | 0.7192 | 0.8021 | -0.2138 |
| Shoot:Root | -0.0897 | -0.0286 | -0.1811 | -0.2699 | 0.7935 |
| Length of longest stem | 0.1255 | 0.2167 | 0.2836 | 0.1056 | 0.3446 |

### Microbial treatments produce unique leaf trait combinations

To determine whether soil microbes induce unique multivariate phenotypes, we first assessed microbial treatment effects on group mean leaf and growth phenotypes. MANOVA found that microbial treatment significantly affected overall leaf traits (F_20, 384□_ = 7.468, P < 0.001) and growth traits (F_16, 432_ = 6.377, P < 0.001; Table S8). These results were confirmed by linear discriminant analysis (Figure 3A-B).

**Figure 3:**
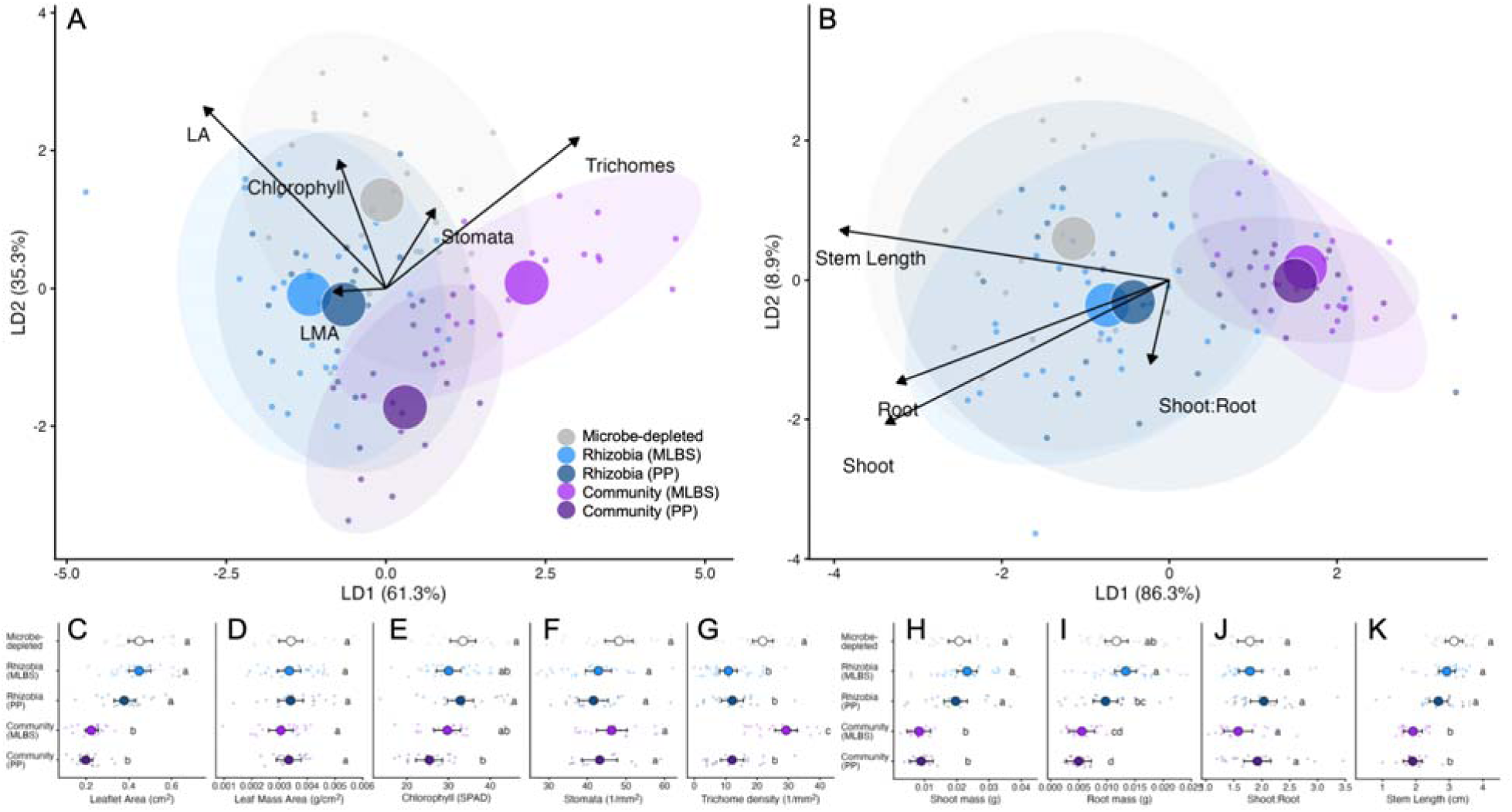
Linear discriminant analysis among microbial treatments for suites of leaf (A) and growth traits (B), with univariate treatment means (C-K). We estimated treatment means and 95% confidence intervals with linear models for leaflet area (C), leaf mass area (D), chlorophyll (E), stomata density (F), trichome density (G), shoot biomass (H), root biomass (I), shoot:root ratio (J), and longest stem length (K). Letters produced by compact letter displays indicate significant differences among treatments for each trait (α = 0.05), with post-hoc tests in Table S4. Treatments sharing a letter are not significantly different.

For leaf traits, LD1 explained 61.3% of treatment differences and was primarily associated with leaf area and trichome density, which loaded oppositely (loadings: −0.713 and 0.755, respectively; Table 2A). LD2 explained an additional 35.3% of the trace, with leaf area (0.658) and trichome density (0.547) again loading most strongly, this time in the same direction. Together, LD1 and LD2 explained 96.6% of treatment differences in leaf traits. Trichome density separated the microbe-depleted and community (MLBS) treatments from the two rhizobia treatments and the community (PP) treatment. Leaflet area primarily distinguished the two rhizobia treatments and the microbe-depleted control from the community treatments. Community treatments had overall reduced leaflet area; the MLBS community had relatively increased trichome density compared to the PP community, which had relatively lower trichome density; Rhizobia treatments both decreased trichome density to the microbe-depleted control with relatively little effect on leaflet area (Figure 3A, C, G). Thus, leaflet area and trichome density jointly drove treatment discrimination, with orthogonal loadings driving unique trait combinations.

**Table 2:** Proportion of trace and structure coefficient trait loadings onto linear discriminant axes for leaf traits (A) and growth traits (B) comparing across all treatment groups. These describe the correlations between the original traits and the axes, and the proportion of group separability explained by each discriminant function.

| A) Leaf traits |  |  |  |  |
| --- | --- | --- | --- | --- |
|  | LD1 | LD2 | LD3 | LD4 |
| **Proportion of trace** | 0.613 | 0.353 | 0.028 | 0.005 |
| Leaf Area | -0.713 | 0.658 | 0.147 | -0.096 |
| Leaf Mass Area | -0.206 | -0.011 | -0.175 | 0.184 |
| Chlorophyll | -0.186 | 0.465 | -0.827 | 0.183 |
| Stomata Density | 0.192 | 0.288 | 0.425 | 0.828 |
| Trichome Density | 0.755 | 0.547 | 0.168 | -0.285 |
| B) Growth traits |  |  |  |  |

|  | LD1 | LD2 | LD3 | LD4 |
| --- | --- | --- | --- | --- |
| **Proportion of trace** | 0.863 | 0.089 | 0.043 | 0.004 |
| Shoot biomass | -0.847 | -0.516 | 0.087 | -0.095 |
| Root biomass | -0.814 | -0.370 | 0.441 | 0.085 |
| Shoot:Root | -0.056 | -0.300 | -0.946 | 0.109 |
| Length of longest stem | -0.983 | 0.179 | -0.018 | -0.042 |

For growth traits, LD1 explained 86.3% of treatment differences and was strongly associated with all size-related traits: stem length (−0.983), shoot biomass (−0.847), and root biomass (−0.814), while shoot:root ratio contributed minimally (−0.056; Table 2B). LD1 primarily separated the two community treatments from the rhizobia and microbe-depleted treatments, reflecting that community-inoculated plants were substantially smaller. LD2 explained only 8.9% of treatment differences, with shoot biomass (−0.516) contributing most strongly, followed by root biomass (−0.370): unlike within-treatment variation, the second discriminant axis did not represent allocation. Together, LD1 and LD2 explained 95.2% of treatment differences in growth traits.

Our global PCA analyses of leaf and growth traits combining all *individuals* in the experiment across treatments yielded similar, but distinct axes of variation compared to the linear discriminant analyses comparing group means. The global first principal component for growth traits explained 64.7% of variation and had strong positive loadings from shoot mass (0.59), root mass (0.59), and the length of the longest stem (0.54), and a weak loading of shoot:root (−0.056). The global first principal component for leaf traits explained 36.3% of variation and had moderately strong positive loadings from leaflet area (0.468), leaf mass area (0.585), chlorophyll content (0.511), moderately strong negative loading from trichome density (−0.421), and weak loading from stomata density, which had a strong negative loading on PC2 (−0.899); leaflet area, leaflet mass area, and trichome density had weak to moderately negative loadings (−0.163, −0.144, −0.378, respectively) (Table S1A). Like the linear discriminant analysis, leaflet area and trichome density loaded orthogonally (oppositely on the first axis of variation, and similarly on the second axis of variation).

Univariate analyses confirmed that leaflet area and trichome density showed the largest and most treatment-specific effects (Figure 3C-K). The community treatments significantly reduced leaflet area compared to the microbe-depleted control (Figure 3C; S2A; Table S4). Trichome density varied substantially among treatments, with contrasting effects: the community (MLBS) treatment had increased trichome density, while the community (PP) treatment and both rhizobia treatments had decreased trichome density compared to the microbe-depleted control (Figure 3K; S2E; Table S4). In contrast, other leaf traits had more muted responses: leaflet mass area did not differ significantly among treatments (p = 0.650), stomata density showed only marginal variation (p = 0.076), and chlorophyll differed primarily because the community (PP) treatment exhibited reduced chlorophyll (Table S3, S4).

For growth traits, shoot biomass, root biomass, and stem length all showed significant treatment effects, with community treatments consistently smaller than rhizobia and microbe-depleted treatments (Figure 3H-K; Table S3). Shoot:root ratio did not differ significantly among treatments (p = 0.097; Figure J; Table S3).

### Microbial effects on nodulation and growth explain leaf area, but not trichome density

We next asked whether variation in growth and rhizobia nodulation explained microbial effects on leaf traits (Figure 4). We interpreted the first principal component of the growth axis—which was strongly correlated with shoot mass, root mass, and the length of the longest stem with similar weights—as an index of general plant performance and growth (Table S1, Figure S1).

**Figure 4:**
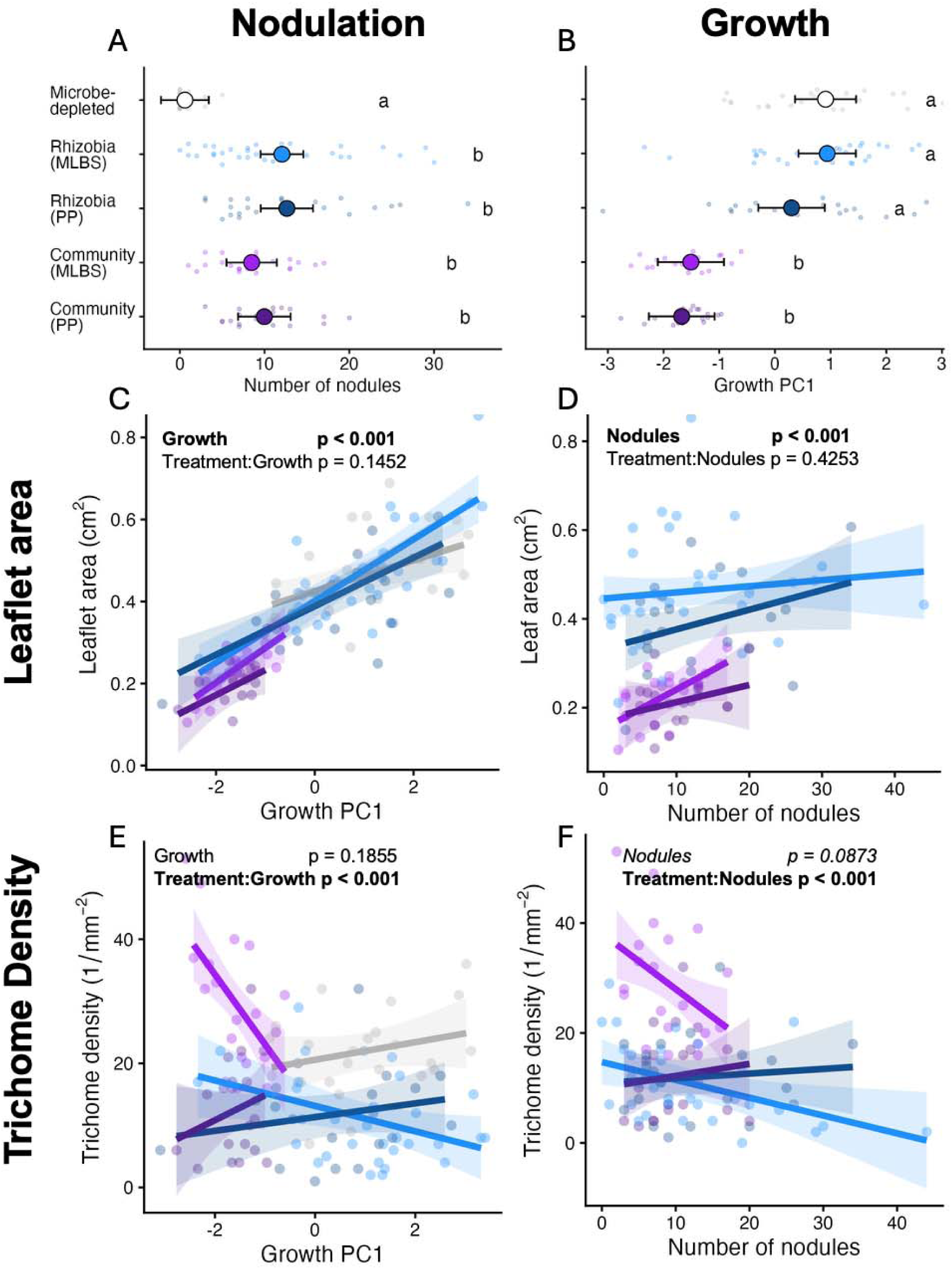
The relationship of focal leaf trait variation with growth and nodulation. Microbes had differential effects on growth (A) and nodulation (B). The relationship between leaflet area and growth PC1 (from global PCA; Table S1) (C) and leaflet area and nodulation (D) were both significantly positive. The relationship between trichome density and growth (E) and nodulation (F) varied depending on treatment. Points are individual plants colored by treatment; ANOVA results are referenced in Table S5, while post-hoc tests for trichome density are in Table S6.

This growth PC1 positively corresponded with leaflet area (p < 0.001; Table S5) without a significant interaction effect by treatment (Figure 4C). Larger plants had larger leaves regardless of microbial treatment, and variation in leaf area was well explained by differences in overall growth. Nodulation also explained variation in leaflet area but not trichome density. Increased nodule number was significantly associated with expanded leaflet area (p = 0.013; Table S5), and this relationship did not depend on treatment (treatment × nodules interaction: p = 0.425; Figure 4D).

On the other hand, nodule number had only a marginally significant—and negatively trending—overall effect on trichome density (p = 0.087), which was treatment-dependent (treatment × nodules interaction: p = 0.040; Table S5; Figure 4F). Trichome density was therefore not strongly or consistently explained by differences in nodulation investment. The growth PC1—our index for general plant performanc—also did not significantly predict trichome density overall (p = 0.186; Table S5). There was a strong treatment × growth interaction (p < 0.001; Figure 4E), driven primarily by the community (MLBS) treatment which had a negative relationship between trichome density and growth which differed from all other treatments (Table S6).

The microbe-depleted treatment showed minimal nodulation, confirming successfully maintained depletion in the field (Figure 4B; Table S3), while inoculated treatments developed an average of 8-12 nodules (Figure S3). Notably, rhizobia treatments did not significantly increase growth compared to the microbe-depleted control, but plants with more nodules were generally larger, suggesting that rhizobia were not exploitative (Figure S3; Table S7).

## Discussion

We found that microbes do not merely shift individual trait means, they also restructure the relationships among traits within an organism. Our results illustrate the extent, as well as the limits, of this restructuring. Soil microbes modified leaf phenotypic integration by altering the covariance structure among traits: microbes mostly increased phenotypic variation, consistently strengthened phenotypic covariances, sometimes reoriented the primary axis of leaf trait covariation, and produced distinct multivariate leaf phenotypes that differed across microbial treatments. Variance in trichome density—a defense trait highly responsive to our microbial treatments—contributed to this microbial restructuring of leaf traits. Compared to leaf traits, growth trait covariance structure was largely robust to microbial manipulation. Together, our findings demonstrate that microbes can modify phenotypic integration, and that a multivariate perspective is necessary to characterize the full extent of these effects.

### Microbes restructured phenotypic covariance in leaf but not growth traits

All four microbial treatments strengthened phenotypic covariances among leaf traits, usually while increasing total phenotypic variation (Figure 2F-G; Table S2). The leaf phenotype was thus overall more integrated for plants that had microbes. Thus we observed that microbes affect both how strongly leaves covary, *and* which traits are most strongly associated, indicating that microbes, like the abiotic environment, can alter trait covariance structure (Schlichting 1989; Callahan and Waller 2000; Wood and Brodie 2015). The direction and magnitude of these effects was not specific to the type of microbial manipulation we applied. Rhizobia and rhizosphere microbial communities had equally strong effects on all measured dimensions of leaf trait integration, although their specific effects differed. This observation suggests that a wide range of microbes, from highly specific endosymbionts to entire microbial communities, have the capacity to impact trait relationships in their hosts.

Growth traits, by contrast, maintained a relatively consistent covariance structure. The dominant axis in all treatments was size, comprising shoot biomass, root biomass, and stem length for all treatments (Table 1, Figure 2F–J). Therefore, growth traits were more robustly integrated than leaf traits for the microbial treatments we manipulated, suggesting that coordinated size scaling may be canalized for early plant growth and is less sensitive to microbial perturbation than leaf traits. We had expected shoot-to-root ratio—an allocation trait—to be especially sensitive to microbial manipulation, since resource-exchange mutualisms and microbial competition might plausibly alter carbon and nitrogen allocation. However, microbial treatments did not significantly alter average shoot:root ratios (Chapin, 1991; Friesen et al. 2011; Figure 2J). Overall, soil microbes mostly shifted plants along a common axis of trait combinations reflecting size, rather than restructuring growth trait architecture itself (Figure 3B).

Our analyses focused on comparisons to the microbe-depleted control. The ecological relevance of this comparison is questionable given that all plants associate with soil microbes in natural conditions (Partida-Martinez and Heil 2011). However, it is important to note that our microbe-depleted control was not a statistical outlier compared to the microbial treatments in any of our nine measured traits (Figure 3C-K). Although our comparisons focused on the microbe-depleted control, there were also qualitative differences among the inoculated microbial treatments themselves. This observation indicates that the microbial effects on phenotypic integration that we documented are ecologically relevant, and that they may extend to heterogeneity in the microbial environment that wild plants may experience (Figure S1).

Phenotypic correlations have often been used as a proxy for underlying genetic relationships between traits that would alter evolutionary trajectories (Cheverud 1982; Roff 1995; Conner and Sterling 1996). If the phenotypic correlations that we documented mirror the genetic relationships among traits, it would mean that microbes could 1) expand evolvability by increasing phenotypic variation, 2) modify the speed of evolution by strengthening or weakening phenotypic correlations, 3) result in novel correlational selection by modifying the direction of phenotypic correlations. However, when and whether phenotypic correlations reflect the underlying genetic correlations remains an open question. Work has also shown that the relationships among traits can differ in strength or even direction across scales of biological organization (Messier et al. 2017; Peiman and Robinson 2017; Agrawal 2020). Therefore, although we can speculate that microbes *might* affect evolutionary potential or evolutionary constraint, given their effects on leaf phenotypic integration, this requires further direct investigation to actually demonstrate.

### Microbial treatments produced distinct combinations of leaf traits, but not growth traits

For the suite of growth traits, there were similar patterns of trait association within and across microbial environments. The same size traits (shoot mass, root mass, and stem length) separated plants in different microbial treatment groups, as well as individuals within and across microbial treatments. The one exception was the rhizosphere microbial community from Pandapas Pond (Figure 2, Figure 3) Thus, although growth consisted of multiple traits, it was a largely one-dimensional phenotype: plants were primarily large or small, which affected roots, shoots, and stem length similarly.

However, differences in leaf traits across microbial treatments did not simply reflect patterns of trait covariance observed within treatments. The traits that separated treatments were not exactly the same as the major axis of phenotypic variation within treatments (Table 1 vs. Table 2). This is consistent with a well documented phenomenon in the plant functional traits literature that patterns of correlation can be eroded or reverse across scales (Donovan et al. 2011b; Messier et al. 2017; Anderegg et al. 2018; Agrawal 2020). This decoupling of within- and between-treatment variation means that knowing how leaf traits covary within any given microbial context does not readily predict which traits or trait combinations will be most strongly affected by microbes overall, across groups.

Our experiment revealed that microbial effects on leaf traits were not strongly aligned with variation along a resource acquisition-conservation trait axis, as predicted by the leaf economic spectrum (LES). The LES is a canonical conceptual framework that describes a fundamental trade-off between resource-acquisitive strategies (characterized by high photosynthetic capacity, high leaf nitrogen, and low leaf mass area) and resource-conservative strategies (lower photosynthetic rates, longer leaf lifespans, and higher leaf mass area) (Donovan et al. 2011b; Reich 2014). Although treatment differences were associated primarily with leaflet area and trichome density, the two were largely decoupled from each other; other LES traits contributed weakly to distinguishing treatment groups (Figure 3A). For all plants across microbial treatments (Figure S1A) and within treatment groups (Figure 2A-E), there was also no strong evidence in our data for a single acquisitive-conservative axis primarily determining leaf traits at either scale: acquisitive traits and conservative traits loaded differently onto multiple dimensions.

While the LES is a powerful framework, the patterns it describes tends to hold up most strongly at large (global, cross-species) scales, with examples of breakdown at smaller scales if insufficient variation is represented (Messier et al. 2017; Anderegg et al. 2018). In addition, there are several reasons that variation in our data would be imperfectly described by the LES. We were missing canonical LES traits like leaf nitrogen content and leaf lifespan, and our measurement of photosynthesis used a SPAD meter to measure relative chlorophyll content rather than measuring photosynthetic rate directly. Additionally, we measured trichome density, a defense trait. Defense traits were not included in the initial formulation of the LES, but they are increasingly becoming integrated into resource economic theory (Agrawal and Fishbein 2006; Vannette and Hunter 2011; Mason and Donovan 2015; Mason et al. 2016). There is evidence that variety in defense traits may lead to certain defense traits (like lignin) being associated with a conservative strategy (Onoda et al. 2011; Kitajima et al. 2013), while other defense types may be inconsistently associated with either axis (Moles et al. 2013), or independently associated with alternative axes of variation (Chauvin et al. 2018).

Nevertheless, we found that microbial effects on defense were a major axis of variation in our data that had no correlation with the growth axis and was independent of the acquisitive-to-conservative spectrum. Trichome defense and leaflet mass area were negatively correlated within treatments, counter to the expectation that both traits should be associated with resource-conservative leaf phenotype (Figure 2A-E). Treatment differences in leaf traits were driven primarily by microbial effects on leaflet area and trichome defense, whose responses were largely decoupled (Figure 3A). Defense and the canonical LES traits loaded oppositely onto the primary axis of variation (representing 36.3% of total variation and 61.3% of variation across groups), perhaps reflecting an alternative axis of variation constituted by a partial tradeoff between growth and defense. However, this alternative axis was not constrained, as plants had unique combinations of leaflet area and trichome density across treatments. Microbial treatments produced distinct multivariate phenotypes which often contradicted expected resource allocation tradeoffs.

### Microbial effects on defense shaped microbial effects on leaf phenotypes, and were not explained by growth or investment in the rhizobia mutualism

Trichome density contributed substantially to observed microbially-mediated changes in leaf covariance structure. Omitting trichome density from the analysis eliminated the microbial effect on total phenotypic variation and substantially reduced differences in the strength of phenotypic correlations across treatments. Thus, while microbial treatments affected trait relationships in multiple ways, trichome density was unusually sensitive to microbial inoculation.

However, microbial effects on trichome density appeared to arise independently from their overall effect on growth (Figure 4). Nor was there a consistent association between rhizobia mutualism investment and trichome density. In contrast, variation in leaflet area—the other leaf trait that was particularly sensitive to microbes— was associated with growth and rhizobia nodulation. This is perhaps not surprising, given that rhizobia provide resources and promote growth (Figure S3), and that leaflet area may be a proxy growth trait, given that it is a measure of size.

The lack of consistent relationship between growth and nodulation trichome density strongly suggests that microbes are more than their stereotyped effects on growth. In fact, different microbial treatments decoupled size and defense, suggesting that they modify growth-defense tradeoffs within treatments (Ku et al. 2024). In fact, only one unique microbial treatment (the MLBS community) exhibited a growth-defense tradeoff. Relative to the microbe-depleted control, the other three treatments affected size and trichome density independently. For example, both community-inoculated treatments were significantly smaller than the microbe-depleted control, but they had opposite effects on trichome density. Together, these results argue against the simplest mechanistic interpretation of microbe-mediated plasticity, in which microbes act primarily as resource providers whose effects cascade through overall plant growth to influence individual traits. Growth-promoting effects clearly influenced some aspects of the leaf phenotype—leaflet area in particular—but trichome density responded to a different signal that varied across microbial types and communities in ways not predicted by growth-level metrics.

### Limitations and Future Directions

The mechanisms by which microbes influence trichome density remain unclear. The decoupling of these effects from growth and nodulation suggests that microbial effects on defense are not mediated simply through resource provisioning. We speculate that microbes might influence trichome density through signaling pathways, such as jasmonic acid-mediated responses, induced systemic resistance, or analogous mechanisms activated by specific microbial taxa rather than generalized resource provisioning (Wagner et al. 2004; Friesen et al. 2011). Yet our experiment did not identify specific microbial taxa in the community treatment, and specific pathways remain untested. Whether microbes disproportionately affect defense across systems—and whether there are consistent mechanisms by which they might modify defense—are open questions.

Our experiment captured early vegetative growth in *M. lupulina*, a relatively short window that may not represent the full trajectory of plant-microbe interactions across the host lifespan. We measured early life traits, which may reflect a sensitive developmental stage when plants invest in the carbon establishment costs of rhizobia nodulation without yet realizing the full payoff of nitrogen benefit (Minchin and Pate 1973; Udvardi and Poole 2013). This may explain why rhizobia did not increase photosynthetic capacity or growth (Figure 3E, H). Whether the effects on phenotypic integration we report here are transient or persist into reproductive maturity, when fitness consequences of altered trait covariance could be measured directly, is unknown. More broadly, the relationship between the **P**-matrix changes we document and the **G**-matrix that constrains evolutionary responses to selection (Cheverud 1988; Wood and Brodie 2015) remains an open and important empirical question. If microbes alter **P** substantially—as our results suggest they can—it is worth asking whether they also alter **G**, and thereby the evolutionary trajectory of host populations that differ in their microbiomes.

If our goal is to understand fitness consequences of microbial effects, field experiments are essential. Hosts are involved in diverse ecological interactions (herbivory, pollination, competition) and experience heterogeneous environmental conditions (shade, drought, resource availability) that shape natural selection. Fitness components cannot be meaningfully assessed in the laboratory, but rigorous microbial manipulation is assumed to require sterile and stringently controlled conditions. We sustained microbial inoculations in the field with relative success, maintaining experimental control in more ecologically realistic and natural conditions. Others have successfully employed similar approaches (Lau and Lennon 2011; Simonsen and Stinchcombe 2014; Petipas et al. 2020), and we encourage more researchers to adopt such field experiments.

Finally, we emphasize the value of multivariate approaches. A multivariate perspective is uniquely powerful to characterize the full scope of microbial effects on host phenotypes. Univariate analyses can identify which traits change and how much, but they cannot reveal how traits are reorganized—whether microbes shift trait means, restructure covariances, or both. We hope that our study encourages more researchers to adopt multivariate frameworks in field-based microbiome experiments.

## References

Agrawal, A. A. 2020. A scale-dependent framework for trade-offs, syndromes, and specialization in organismal biology. Ecology 101:e02924.

Agrawal, A. A., and M. Fishbein. 2006. Plant defense syndromes. Ecology 87:S132–149.

Anderegg, L. D. L., L. T. Berner, G. Badgley, M. L. Sethi, B. E. Law, and J. HilleRisLambers. 2018. Within-species patterns challenge our understanding of the leaf economics spectrum. (J. Penuelas, ed.)Ecology Letters 21:734–744.

Arnold, S. J., R. Bürger, P. A. Hohenlohe, B. C. Ajie, and A. G. Jones. 2008. Understanding the Evolution and Stability of the G-Matrix. Evolution; international journal of organic evolution 62:2451–2461.

Bogar, L. M. 2024. Modified source–sink dynamics govern resource exchange in ectomycorrhizal symbiosis. New Phytologist 242:1523–1528.

Brooks, M. E. K., Kristensen, K., van Benthem, J. A., Magnusson, C., Berg W. A., Nielsen, H., Skaug J., et al. 2017. glmmTMB Balances Speed and Flexibility Among Packages for Zero-inflated Generalized Linear Mixed Modeling. The R Journal 9:378.

Callahan, H. S., and D. M. Waller. 2000. Phenotypic Integration and the Plasticity of Integration in an Amphicarpic Annual. International Journal of Plant Sciences 161:89–98.

Chapin, F. S. 1991. Integrated Responses of Plants to Stress. BioScience 41:29–36.

Chauvin, K. M., G. P. Asner, R. E. Martin, W. J. Kress, S. J. Wright, and C. B. Field. 2018. Decoupled dimensions of leaf economic and anti-herbivore defense strategies in a tropical canopy tree community. Oecologia 186:765–782.

Chenoweth, S. F., H. D. Rundle, and M. W. Blows. 2010. The Contribution of Selection and Genetic Constraints to Phenotypic Divergence. The American Naturalist 175:186–196.

Cheverud, J. M. 1982. PHENOTYPIC, GENETIC, AND ENVIRONMENTAL MORPHOLOGICAL INTEGRATION IN THE CRANIUM. Evolution 36:499–516.

Cheverud, J. M. 1988. A COMPARISON OF GENETIC AND PHENOTYPIC CORRELATIONS. Evolution; International Journal of Organic Evolution 42:958–968.

Conner, J. K., and A. Sterling. 1996. Selection for independence of floral and vegetative traits: evidence from correlation patterns in five species. Canadian Journal of Botany 74:642–644.

Damián, X., S. Ochoa-López, A. Gaxiola, J. Fornoni, C. A. Domínguez, and K. Boege. 2020. Natural selection acting on integrated phenotypes: covariance among functional leaf traits increases plant fitness. New Phytologist 225:546–557.

Day, R. W., and G. P. Quinn. 1989. Comparisons of Treatments After an Analysis of Variance in Ecology. Ecological Monographs 59:433–463.

De Jong, G. 1993. Covariances Between Traits Deriving From Successive Allocations of a Resource. Functional Ecology 7:75.

Delahaie, B., A. Charmantier, S. Chantepie, D. Garant, M. Porlier, and C. Teplitsky. 2017. Conserved G-matrices of morphological and life-history traits among continental and island blue tit populations. Heredity 119:76–87.

Díaz, S., J. Kattge, J. H. C. Cornelissen, I. J. Wright, S. Lavorel, S. Dray, B. Reu, et al. 2016. The global spectrum of plant form and function. Nature 529:167–171.

Donovan, L. A., H. Maherali, C. M. Caruso, H. Huber, and H. De Kroon. 2011a. The evolution of the worldwide leaf economics spectrum. Trends in Ecology & Evolution 26:88–95.

Donovan, L. A., H. Maherali, C. M. Caruso, H. Huber, and H. de Kroon. 2011b. The evolution of the worldwide leaf economics spectrum. Trends in Ecology & Evolution 26:88–95.

Douglas, A. E. n.d. Aphids and Their Symbiotic Bacteria.

Dunnett, C. W. 1955. A Multiple Comparison Procedure for Comparing Several Treatments with a Control. Journal of the American Statistical Association 50:1096–1121.

Finch, H. 2005. Comparison of the Performance of Nonparametric and Parametric MANOVA Test Statistics when Assumptions Are Violated. Methodology 1:27–38.

Fontaine, S. S., P. M. Mineo, and K. D. Kohl. 2022. Experimental manipulation of microbiota reduces host thermal tolerance and fitness under heat stress in a vertebrate ectotherm. Nature Ecology & Evolution 6:405–417.

Fox, J., and S. Weisberg. 2019. An R Companion to Applied Regression (3rd ed.). Sage, Thousand Oaks, CA.

Friendly, M., & Fox, J. (2025). candisc: Visualizing Generalized Canonical Discriminant and Canonical Correlation Analysis (R package version 1.0.0).

Friesen, M. L., S. S. Porter, S. C. Stark, E. J. von Wettberg, J. L. Sachs, and E. Martinez-Romero. 2011. Microbially Mediated Plant Functional Traits. Annual Review of Ecology, Evolution, and Systematics 42:23–46.

Gano-Cohen, K. A., C. E. Wendlandt, K. A. Moussawi, P. J. Stokes, K. W. Quides, A. J. Weisberg, J. H. Chang, et al. n.d. Recurrent mutualism breakdown events in a legume rhizobia metapopulation 9.

Hansen, T. F., C. Pélabon, and D. Houle. 2011. Heritability is not Evolvability. Evolutionary Biology 38:258–277.

Harrison, T. L., C. W. Wood, I. L. Borges, and J. R. Stinchcombe. 2017a. No evidence for adaptation to local rhizobial mutualists in the legume Medicago lupulina. Ecology and Evolution 7:4367–4376.

Harrison, T. L., C. W. Wood, K. D. Heath, and J. R. Stinchcombe. 2017b. Geographically structured genetic variation in the Medicago lupulina–Ensifer mutualism. Evolution 71:1787–1801.

Hartig F (2026). DHARMa: Residual Diagnostics for Hierarchical (Multi-Level / Mixed) Regression Models. R package version 0.5.0, https://github.com/florianhartig/dharma.

Hawkes, C. V., R. Kjøller, J. M. Raaijmakers, L. Riber, S. Christensen, S. Rasmussen, J. H. Christensen, et al. 2021. Extension of Plant Phenotypes by the Foliar Microbiome. Annual Review of Plant Biology 72:823–846.

Heath, K. D., and J. R. Stinchcombe. 2014. EXPLAINING MUTUALISM VARIATION: A NEW EVOLUTIONARY PARADOX? Evolution 68:309–317.

Heath-Heckman, E. A. C., A. A. Gillette, R. Augustin, M. X. Gillette, W. E. Goldman, and M. J. McFall-Ngai. 2014. Shaping the microenvironment: evidence for the influence of a host galaxin on symbiont acquisition and maintenance in the squid-Vibrio symbiosis. Environmental Microbiology 16:3669–3682.

Higashi, C. H. V., V. Patel, B. Kamalaker, R. Inaganti, A. Bressan, J. A. Russell, and K. M. Oliver. 2024. Another tool in the toolbox: Aphid-specific Wolbachia protect against fungal pathogens. Environmental Microbiology 26:e70005.

Houle, D., C. Pélabon, G. Wagner, and T. Hansen. 2011. Measurement and Meaning in Biology. The Quarterly review of biology 86:3–34.

Jung, S. C., A. Martinez-Medina, J. A. Lopez-Raez, and M. J. Pozo. 2012. Mycorrhiza-Induced Resistance and Priming of Plant Defenses. Journal of Chemical Ecology 38:651–664.

Kitajima, K., R. A. Cordero, and S. J. Wright. 2013. Leaf life span spectrum of tropical woody seedlings: effects of light and ontogeny and consequences for survival. Annals of Botany 112:685–699.

Ku, Y., Y. Liao, S. Chiou, H. Lam, and C. Chan. 2024. From trade-off to synergy: microbial insights into enhancing plant growth and immunity. Plant Biotechnology Journal 22:2461–2471.

Lande, R., and S. J. Arnold. 1983. The Measurement of Selection on Correlated Characters. Evolution 37:1210–1226.

Lange, C., S. Boyer, T. M. Bezemer, M.-C. Lefort, M. K. Dhami, E. Biggs, R. Groenteman, et al. 2023. Impact of intraspecific variation in insect microbiomes on host phenotype and evolution. The ISME Journal.

Lau, J. A., and J. T. Lennon. 2011. Evolutionary ecology of plant–microbe interactions: soil microbial structure alters selection on plant traits. New Phytologist 192:215–224.

Lenth R, Piaskowski J (2026). emmeans: Estimated Marginal Means, aka Least-Squares Means. R package.

Ling, N., T. Wang, and Y. Kuzyakov. 2022. Rhizosphere bacteriome structure and functions. Nature Communications 13:836.

Martin, A. R., R. O. Mariani, P. Dörr de Quadros, and R. R. Fulthorpe. 2022. The influence of biofertilizers on leaf economics spectrum traits in a herbaceous crop. Journal of Experimental Botany 73:7552–7563.

Mason, C. M., A. W. Bowsher, B. L. Crowell, R. M. Celoy, C.-J. Tsai, and L. A. Donovan. 2016. Macroevolution of leaf defenses and secondary metabolites across the genus Helianthus. New Phytologist 209:1720–1733.

Mason, C. M., and L. A. Donovan. 2015. Does investment in leaf defenses drive changes in leaf economic strategy? A focus on whole-plant ontogeny. Oecologia 177:1053–1066.

Masson-Boivin, C., and J. L. Sachs. 2018. Symbiotic nitrogen fixation by rhizobia-the roots of a success story. Current Opinion in Plant Biology 44:7–15.

McGlothlin, J. W., M. E. Kobiela, H. V. Wright, J. J. Kolbe, J. B. Losos, and E. D. Brodie. 2022. Conservation and Convergence of Genetic Architecture in the Adaptive Radiation of Anolis Lizards. The American Naturalist 200:E207–E220.

Messier, J., B. J. McGill, B. J. Enquist, and M. J. Lechowicz. 2017. Trait variation and integration across scales: is the leaf economic spectrum present at local scales? Ecography 40:685–697.

Midway, S., M. Robertson, S. Flinn, and M. Kaller. 2020. Comparing multiple comparisons: practical guidance for choosing the best multiple comparisons test. PeerJ 8:e10387.

Minchin, F. R., and J. S. Pate. 1973. The Carbon Balance of a Legume and the Functional Economy of its Root Nodules. Journal of Experimental Botany 24:259–271.

Moles, A. T., B. Peco, I. R. Wallis, W. J. Foley, A. G. B. Poore, E. W. Seabloom, P. A. Vesk, et al. 2013. Correlations between physical and chemical defences in plants: tradeoffs, syndromes, or just many different ways to skin a herbivorous cat? New Phytologist 198:252–263.

Murren, C. J. 2002. Phenotypic integration in plants. Plant Species Biology 17:89–99.

Mylona, P., K. Pawlowski, and T. Bisseling’. 1995. Symbiotic Nitrogen Fixation. The Plant Cell 7:869–885.

O’Brien, A. M., N. A. Ginnan, M. Rebolleda-Gómez, and M. R. Wagner. 2021. Microbial effects on plant phenology and fitness. American Journal of Botany 108:1824–1837.

O’Brien, A. M., R. J. H. Sawers, S. Y. Strauss, and J. Ross-Ibarra. 2019. Adaptive phenotypic divergence in an annual grass differs across biotic contexts*. Evolution 73:2230–2246.

Olson, C. L. 1976. On choosing a test statistic in multivariate analysis of variance. Psychological Bulletin 83:579–586.

Onoda, Y., M. Westoby, P. B. Adler, A. M. F. Choong, F. J. Clissold, J. H. C. Cornelissen, S. Díaz, et al. 2011. Global patterns of leaf mechanical properties. Ecology Letters 14:301–312.

Partida-Martinez, L. P. P., and M. Heil. 2011. The Microbe-Free Plant: Fact or Artifact? Frontiers in Plant Science 2.

Peiman, K. S., and B. W. Robinson. 2017. Comparative Analyses of Phenotypic Trait Covariation within and among Populations. The American Naturalist 190:451–468.

Petipas, R. H., A. C. Wruck, and M. A. Geber. 2020. Microbe-mediated local adaptation to limestone barrens is context dependent. Ecology 101:e03092.

Pigliucci, M. 2003. Phenotypic integration: studying the ecology and evolution of complex phenotypes. Ecology Letters 6:265–272.

Prashar, P., N. Kapoor, and S. Sachdeva. 2014. Rhizosphere: its structure, bacterial diversity and significance. Reviews in Environmental Science and Bio/Technology 13:63–77.

Pringle, E. G. 2016. Integrating plant carbon dynamics with mutualism ecology. New Phytologist 210:71–75.

R Core Team (2026). R: A Language and Environment for Statistical Computing. R Foundation for Statistical Computing, Vienna, Austria. R Project

Rahman, A., M. Manci, C. Nadon, I. A. Perez, W. F. Farsamin, M. T. Lampe, T. H. Le, et al. 2023. Competitive interference among rhizobia reduces benefits to hosts. Current Biology 33:2988–3001.e4.

Reich, P. B. 2014. The world-wide ‘fast–slow’ plant economics spectrum: a traits manifesto. Journal of Ecology 102:275–301.

Riley, A. B., M. A. Grillo, B. Epstein, P. Tiffin, and K. D. Heath. 2023. Discordant population structure among rhizobium divided genomes and their legume hosts. Molecular Ecology 32:2646–2659.

Roff, D. A. 1995. The estimation of genetic correlations from phenotypic correlations: a test of Cheverud’s conjecture. Heredity 74:481–490.

Sachs, J. L., S. W. Kembel, A. H. Lau, and E. L. Simms. 2009. In Situ Phylogenetic Structure and Diversity of Wild Bradyrhizobium Communities. Applied and Environmental Microbiology 75:4727–4735.

Schlichting, C. D. 1989. Phenotypic Integration and Environmental Change. BioScience 39:460–464.

Schluter, D. 1996. ADAPTIVE RADIATION ALONG GENETIC LINES OF LEAST RESISTANCE. Evolution; International Journal of Organic Evolution 50:1766–1774.

Schneider, C. A., W. S. Rasband, and K. W. Eliceiri. 2012. NIH Image to ImageJ: 25 years of image analysis. Nature Methods 9:671–675.

Sgrò, C. M., and A. A. Hoffmann. 2004. Genetic correlations, tradeoffs and environmental variation. Heredity 93:241–248.

Simonsen, A. K., and J. R. Stinchcombe. 2014. Herbivory eliminates fitness costs of mutualism exploiters. New Phytologist 202:651–661.

Turkington, R., and P. B. Cavers. 1979. THE BIOLOGY OF CANADIAN WEEDS.: 33. Medicago lupulina L. Canadian Journal of Plant Science 59:99–110.

Udvardi, M., and P. S. Poole. 2013. Transport and Metabolism in Legume-Rhizobia Symbioses. Annual Review of Plant Biology 64:781–805.

Vannette, R. L., and M. D. Hunter. 2011. Plant defence theory re-examined: nonlinear expectations based on the costs and benefits of resource mutualisms. Journal of Ecology 99:66–76.

Venables, W. N., & Ripley, B. D. (2002). Modern Applied Statistics with S (4th ed.). Springer.(Alternatively, you can cite Ripley, B. D. (1996). Pattern Recognition and Neural Networks. Cambridge University Press).

Vessey, J. K. 2003. Plant growth promoting rhizobacteria as biofertilizers. Plant and Soil 255:571–586.

Vorholt, J. A., C. Vogel, C. I. Carlström, and D. B. Müller. 2017. Establishing Causality: Opportunities of Synthetic Communities for Plant Microbiome Research. Cell Host & Microbe 22:142–155.

Wagner, G. J., E. Wang, and R. W. Shepherd. 2004. New Approaches for Studying and Exploiting an Old Protuberance, the Plant Trichome. Annals of Botany 93:3–11.

Wagner, M. R., D. S. Lundberg, D. Coleman-Derr, S. G. Tringe, J. L. Dangl, and T. Mitchell-Olds. 2014. Natural soil microbes alter flowering phenology and the intensity of selection on flowering time in a wild Arabidopsis relative. (J. Klironomos, ed.)Ecology Letters 17:717–726.

Wickham, H. (2016). ggplot2: Elegant Graphics for Data Analysis. Springer-Verlag New York. ISBN 978–3-319-24277-4.

Wood, C. W., and E. D. Brodie. 2015. Environmental effects on the structure of the G-matrix: ENVIRONMENTAL EFFECTS ON GENETIC CORRELATIONS. Evolution 69:2927–2940.

Wood, C. W., M. M. Howard, E. Yi, K. D. Heath, K. R. O’Keeffe, and J. A. Lau. 2026. Integrating nutritional mutualists into the evolution of defense. Trends in Ecology & Evolution S0169534726001096.

Zhao, H., H. Liao, A. Biere, and S. Peng. 2026. Plant economics traits predict plant carbon allocation and responsiveness to arbuscular mycorrhizal fungi under varying precipitation. Functional Ecology 40:1921–1932.

