## Supplemental Figures and Tables for "Strong effects of soil microbes on early-life defense alter phenotypic correlations in plant leaves"

**Figure S1: Global PCA of leaf (A) and growth (B) traits on the full phenotypic variation expressed across all microbial treatments.** Trait loadings are represented with labeled arrows.

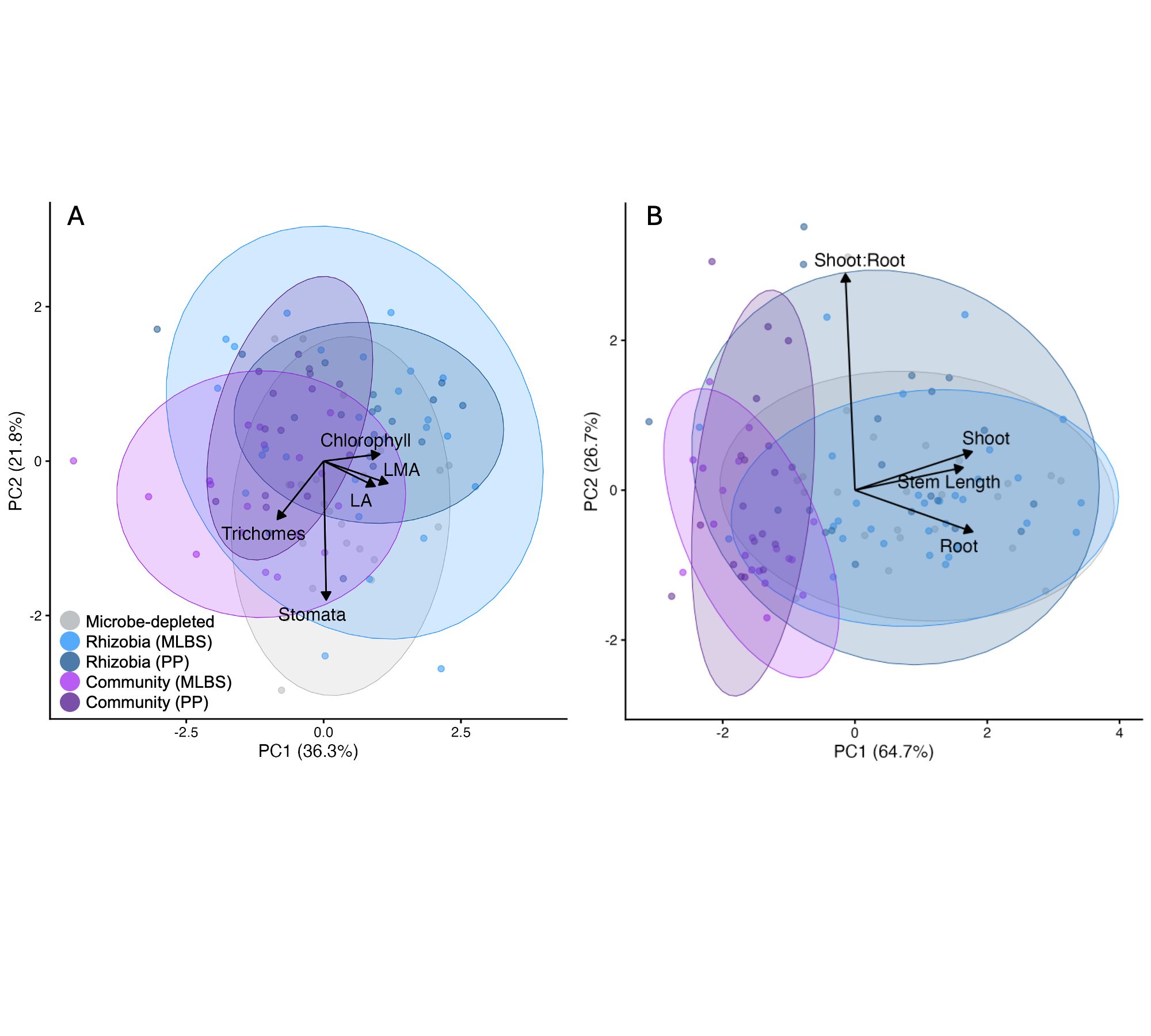

**Table S1: The proportion of variance and trait loadings for global principal components analysis of leaf traits (A) and growth traits (B), conducted on full phenotypic variation expressed across all microbial treatments.** The PC1 of the growth analysis was used as the composite measure in Figure 4 to test whether overall growth and performance drove variation in leaflet area or trichome density.

| A) **Leaf traits** | | | | | |
| --- | --- | --- | --- | --- | --- |
|  | **PC1** | **PC2** | **PC3** | **PC4** | **PC5** |
| *Proportion of Variance* | *0.3628* | *0.2176* | *0.1904* | *0.1432* | *0.086* |
| Leaf Area | 0.4682 | -0.1633 | 0.4073 | -0.7536 | 0.1422 |
| Leaf Mass Area | 0.5845 | -0.1438 | -0.3036 | 0.3526 | 0.649 |
| Chlorophyll | 0.5112 | 0.0454 | 0.5125 | 0.4943 | -0.4791 |
| Stomata Density | 0.0241 | -0.8989 | -0.2677 | -0.0002 | -0.346 |
| Trichome Density | -0.421 | -0.3775 | 0.6384 | 0.2516 | 0.4575 |
| B) **Growth traits** | | | | | |
|  | **PC1** | **PC2** | **PC3** | **PC4** |  |
| *Proportion of Variance* | *0.6474* | *0.2673* | *0.0795* | *0.0057* |  |
| Shoot biomass | 0.5907 | 0.1715 | 0.4144 | 0.6708 |  |
| Root biomass | 0.5932 | -0.1849 | 0.3586 | -0.6966 |  |
| Shoot:Root | -0.0483 | 0.9624 | 0.0822 | -0.2543 |  |
| Length of longest stem | 0.5448 | 0.1007 | -0.8324 | 0.0087 |  |

**Table S2: Pairwise matrix comparison values with permutation test boundaries for leaf traits (A) and growth traits (B)** (corresponding values from Figure 2K-P)**.** The permutation ranges provide significance thresholds to test whether microbial treatments altered trait covariance structure.

| **Leaf traits** | | | | |
| --- | --- | --- | --- | --- |
|  | Rhizobia (MLBS) | Rhizobia (PP) | Community (MLBS) | Community (PP) |
| Volume | 0.9266 | 0.7669 | 0.226 | 0.6196 |
| *Volume (95% CI)* | -0.4871 — 0.4793 | -0.604 — 0.5966 | -0.5542 — 0.5693 | -0.5822 — 0.5979 |
| Proportion of Pmax | 0.3153 | 0.302 | 0.2895 | 0.374 |
| *Proportion of Pmax (95% CI)* | -0.216 — 0.2226 | -0.215 — 0.2119 | -0.2349 — 0.2198 | -0.2401 — 0.2441 |
| Angle between Pmax | 36.85 | 21.87 | 63.77 | 20.87 |
| *Angle between Pmax (95% CI)* | 26.66 | 31.82 | 46.44 | 26.23 |
| **Growth traits** | | | | |
|  | Rhizobia (MLBS) | Rhizobia (PP) | Community (MLBS) | Community (PP) |
| Volume | -0.1145 | 0.1424 | -0.04806 | -0.3971 |
| *Volume (95% CI)* | -0.6034 — 0.5909 | -0.607 — 0.6354 | -0.5196-0.5494 | -0.4975 — 0.5182 |
| Proportion of Pmax | 0.004783 | -0.04792 | -0.01542 | -0.2129 |
| *Proportion of Pmax (95% CI)* | -0.173 — 0.1715 | -0.1611 — 0.1552 | -0.1335-0.1485 | -0.1197 — 0.1209 |
| Angle between Pmax | 7.574 | 10.9 | **13.44** | **83.58** |
| *Angle between Pmax (95% CI)* | 20.63 | 26.35 | 12.57 | 15.32 |

**Figure S2: Cohen’s d effect sizes for leaf and growth traits.**

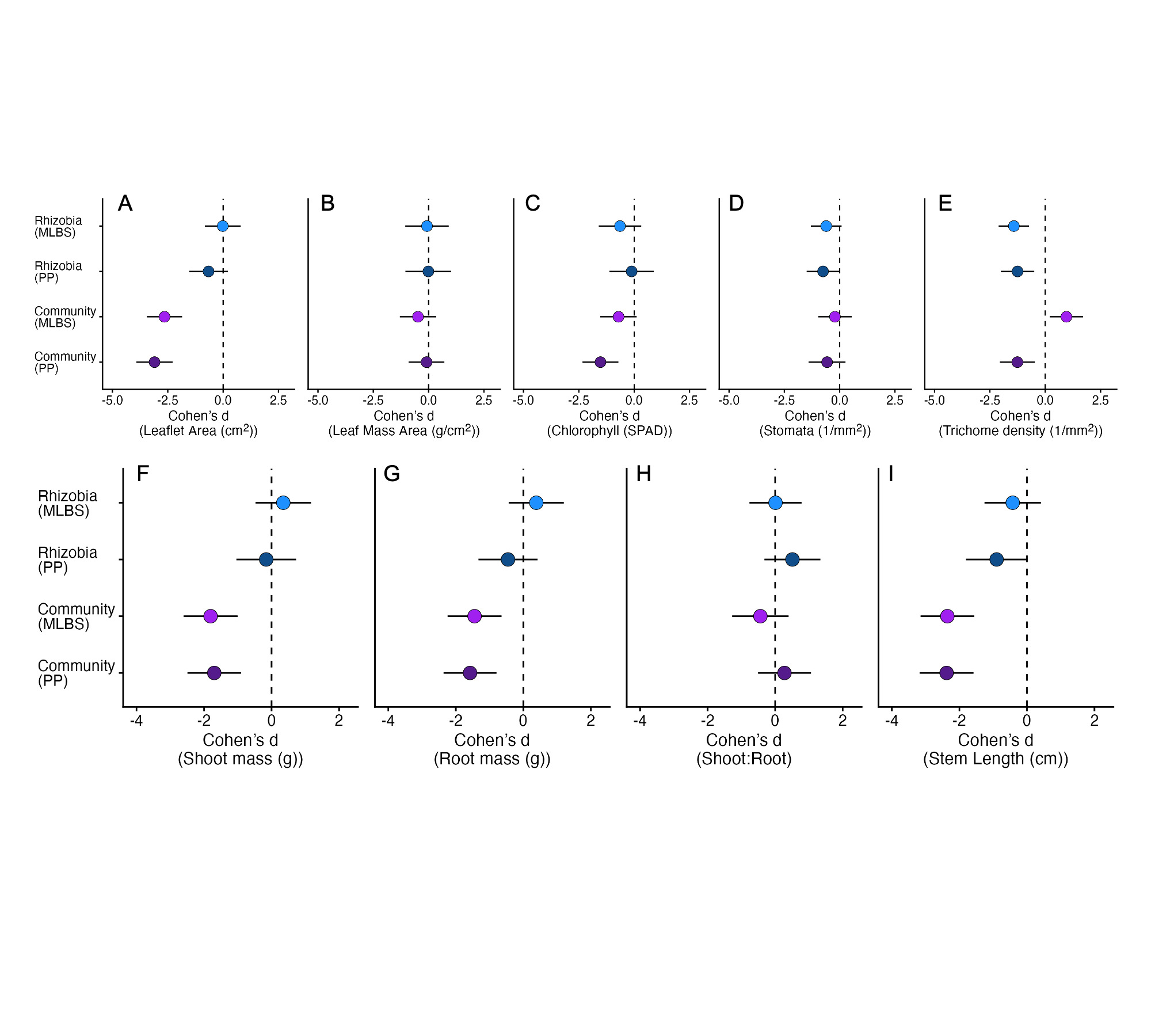

**Table S3: ANOVA results, estimated treatment means, and effect sizes compared to the microbe-depleted control for each leaf, growth, and nodule trait.** F-statistics, degrees of freedom, p-values, estimated marginal means, and Cohen’s d effect sizes for all traits. Corresponds to Figure 3C-K.

| A. Anova differences between leaf, growth, and nodule count traits by treatment. | | | | | | |
| --- | --- | --- | --- | --- | --- | --- |
|  |  |  |  | **Chisq** | **Df** | **P value** |
| **Leaf traits** | | **Leaflet Area** | | 156.477 | 4 | **<0.001** |
|  |  | Leaflet Mass Area | | 2.468 | 4 | 0.650 |
|  |  | **Chlorophyll** | | 26.663 | 4 | **<0.001** |
|  |  | *Stomata* | | *8.461* | *4* | *0.076* |
|  |  | **Trichome density** | | 88.317 | 4 | **<0.001** |
| **Growth traits** | | **Shoot mass** | | 69.946 | 4 | **<0.001** |
|  |  | **Root mass** | | 55.853 | 4 | **<0.001** |
|  |  | *Shoot:Root* | | 7.85 | 4 | *0.0972* |
|  |  | **Stem Length** | | 93.89 | 4 | **<0.001** |
|  | | **Nodule count** | | 44.1 | 4 | **<0.001** |
| B. Estimated means and effect sizes (with 95% CIs) of leaf, growth, and nodule count traits. | | | | | | |
|  |  | **TMD** | **TRA** | **TRB** | **TCM** | **TCP** |
| **Leaf traits** | **Leaflet Area** | | | | | |
|  | Estimated Mean | 0.45 | 0.448 | 0.378 | 0.224 | 0.199 |
|  | *emmeans 95% CI* | *0.396 — 0.51* | *0.4 — 0.502* | *0.331 — 0.432* | *0.196 — 0.256* | *0.171 — 0.231* |
|  | Cohen's d (effect size) | NA | -0.014 | -0.658 | -2.648 | -3.098 |
|  | 95% CI range | NA | -0.823 — 0.795 | -1.533 — 0.216 | -3.444 — -1.852 | -3.921 — -2.275 |
|  | **Leaflet Mass Area** | | | | | |
|  | Estimated Mean | 0.003 | 0.003 | 0.003 | 0.003 | 0.003 |
|  | *emmeans 95% CI* | *0.003 — 0.004* | *0.003 — 0.004* | *0.003 — 0.004* | *0.003 — 0.003* | *0.003 — 0.004* |
|  | Cohen's d (effect size) | NA | -0.073 | -0.013 | -0.481 | -0.094 |
|  | 95% CI range | NA | -1.055 — 0.908 | -1.047 — 1.022 | -1.303 — 0.34 | -0.903 — 0.715 |
|  | **Chlorophyll** | | | | | |
|  | Estimated Mean | 33.467 | 30.06 | 32.861 | 29.704 | 25.349 |
|  | *emmeans 95% CI* | *30.392 — 36.542* | *27.146 — 32.973* | *29.691 — 36.031* | *26.543 — 32.865* | *22.171 — 28.528* |
|  | Cohen's d (effect size) | NA | -0.639 | -0.114 | -0.706 | -1.523 |
|  | 95% CI range | NA | -1.6 — 0.322 | -1.118 — 0.89 | -1.527 — 0.115 | -2.332 — -0.713 |
|  | **Stomata** | | | | | |
|  | Estimated Mean | 48.196 | 42.833 | 41.571 | 46.316 | 43.179 |
|  | *emmeans 95% CI* | *44.597 — 51.794* | *39.512 — 46.155* | *37.805 — 45.338* | *42.356 — 50.275* | *38.566 — 47.791* |
|  | Cohen's d (effect size) | NA | -0.609 | -0.752 | -0.213 | -0.57 |
|  | 95% CI range | NA | -1.305 — 0.087 | -1.493 — -0.012 | -0.974 — 0.547 | -1.401 — 0.262 |
|  | **Trichome density** | | | | | |
|  | Estimated Mean | 21.808 | 10.842 | 12.124 | 29.237 | 12.065 |
|  | *emmeans 95% CI* | *18.58 — 25.036* | *7.994 — 13.689* | *8.63 — 15.617* | *25.687 — 32.787* | *8.407 — 15.723* |
|  | Cohen's d (effect size) | NA | -1.415 | -1.25 | 0.959 | -1.257 |
|  | 95% CI range | NA | -2.102 — -0.728 | -2.004 — -0.496 | 0.205 — 1.712 | -2.046 — -0.468 |
| **Growth traits** | **Shoot mass** | | | | | |
|  | Estimated Mean | 0.021 | 0.023 | 0.02 | 0.008 | 0.009 |
|  | *emmeans 95% CI* | *0.017 — 0.024* | *0.02 — 0.026* | *0.016 — 0.023* | *0.005 — 0.012* | *0.005 — 0.013* |
|  | Cohen's d (effect size) | NA | 0.348 | -0.159 | -1.805 | -1.698 |
|  | 95% CI range | NA | -0.473 — 1.168 | -1.044 — 0.726 | -2.606 — -1.004 | -2.492 — -0.903 |
|  | **Root mass** | | | | | |
|  | Estimated Mean | 0.012 | 0.013 | 0.01 | 0.006 | 0.005 |
|  | *emmeans 95% CI* | *0.01 — 0.014* | *0.011 — 0.015* | *0.008 — 0.012* | *0.003 — 0.008* | *0.003 — 0.007* |
|  | Cohen's d (effect size) | NA | 0.383 | -0.455 | -1.445 | -1.578 |
|  | 95% CI range | NA | -0.433 — 1.199 | -1.33 — 0.42 | -2.244 — -0.646 | -2.362 — -0.794 |
|  | **Shoot:Root** | | | | | |
|  | Estimated Mean | 1.787 | 1.792 | 2.033 | 1.576 | 1.92 |
|  | *emmeans 95% CI* | *1.567 — 2.007* | *1.586 — 1.999* | *1.798 — 2.269* | *1.319 — 1.833* | *1.679 — 2.161* |
|  | Cohen's d (effect size) | NA | 0.011 | 0.512 | -0.437 | 0.277 |
|  | 95% CI range | NA | -0.761 — 0.784 | -0.317 — 1.341 | -1.274 — 0.399 | -0.508 — 1.061 |
|  | **Stem Length** | | | | | |
|  | Estimated Mean | 3.139 | 2.92 | 2.672 | 1.912 | 1.901 |
|  | *emmeans 95% CI* | *2.881 — 3.397* | *2.681 — 3.159* | *2.396 — 2.948* | *1.636 — 2.188* | *1.618 — 2.183* |
|  | Cohen's d (effect size) | NA | -0.421 | -0.898 | -2.357 | -2.379 |
|  | 95% CI range | NA | -1.258 — 0.416 | -1.806 — 0.01 | -3.151 — -1.563 | -3.177 — -1.581 |
|  | **Nodule count** | | | | | |
|  | Estimated Mean | 0.592 | 12.045 | 12.628 | 8.472 | 9.968 |
|  | *emmeans 95% CI* | -2.231 — 3.414 | 9.521 — 14.569 | 9.544 — 15.711 | 5.506 — 11.437 | 6.869 — 13.067 |

**Table S4: Posthoc tests for each of these univariate leaf/growth traits**

|  | Contrast | Estimate | SE | z.ratio | P value |
| --- | --- | --- | --- | --- | --- |
| Leaflet Area | Microbe-depleted - Rhizobia (MLBS) | 0.004 | 0.087 | 0.043 | 1.000 |
|  | Microbe-depleted - Rhizobia (PP) | 0.173 | 0.094 | 1.846 | 0.347 |
|  | Microbe-depleted - Community (MLBS) | 0.697 | 0.085 | 8.159 | <0.001 |
|  | Microbe-depleted - Community (PP) | 0.816 | 0.088 | 9.231 | <0.001 |
|  | Rhizobia (MLBS) - Rhizobia (PP) | 0.170 | 0.077 | 2.211 | 0.176 |
|  | Rhizobia (MLBS) - Community (MLBS) | 0.694 | 0.087 | 7.976 | <0.001 |
|  | Rhizobia (MLBS) - Community (PP) | 0.812 | 0.098 | 8.326 | <0.001 |
|  | Rhizobia (PP) - Community (MLBS) | 0.524 | 0.095 | 5.510 | <0.001 |
|  | Rhizobia (PP) - Community (PP) | 0.642 | 0.103 | 6.212 | <0.001 |
|  | Community (MLBS) - Community (PP) | 0.119 | 0.100 | 1.189 | 0.758 |
| Leaflet Mass Area | Contrast | Estimate | SE | z.ratio | P value |
|  | Microbe-depleted - Rhizobia (MLBS) | 0.000 | 0.000 | 0.183 | 1.000 |
|  | Microbe-depleted - Rhizobia (PP) | 0.000 | 0.000 | 0.030 | 1.000 |
|  | Microbe-depleted - Community (MLBS) | 0.000 | 0.000 | 1.437 | 0.604 |
|  | Microbe-depleted - Community (PP) | 0.000 | 0.000 | 0.285 | 0.999 |
|  | Rhizobia (MLBS) - Rhizobia (PP) | 0.000 | 0.000 | -0.202 | 1.000 |
|  | Rhizobia (MLBS) - Community (MLBS) | 0.000 | 0.000 | 1.052 | 0.831 |
|  | Rhizobia (MLBS) - Community (PP) | 0.000 | 0.000 | 0.052 | 1.000 |
|  | Rhizobia (PP) - Community (MLBS) | 0.000 | 0.000 | 1.120 | 0.796 |
|  | Rhizobia (PP) - Community (PP) | 0.000 | 0.000 | 0.197 | 1.000 |
|  | Community (MLBS) - Community (PP) | 0.000 | 0.000 | -1.054 | 0.830 |
| Chlorophyll | Contrast | Estimate | SE | z.ratio | P value |
|  | Microbe-depleted - Rhizobia (MLBS) | 3.407 | 2.088 | 1.632 | 0.477 |
|  | Microbe-depleted - Rhizobia (PP) | 0.606 | 2.182 | 0.278 | 0.999 |
|  | Microbe-depleted - Community (MLBS) | 3.763 | 1.784 | 2.109 | 0.216 |
|  | Microbe-depleted - Community (PP) | 8.117 | 1.759 | 4.615 | **<0.001** |
|  | Rhizobia (MLBS) - Rhizobia (PP) | -2.801 | 1.561 | -1.794 | 0.377 |
|  | Rhizobia (MLBS) - Community (MLBS) | 0.356 | 2.033 | 0.175 | 1.000 |
|  | Rhizobia (MLBS) - Community (PP) | 4.710 | 2.110 | 2.232 | 0.168 |
|  | Rhizobia (PP) - Community (MLBS) | 3.157 | 2.181 | 1.448 | 0.597 |
|  | Rhizobia (PP) - Community (PP) | 7.511 | 2.159 | 3.479 | **0.00458** |
|  | Community (MLBS) - Community (PP) | 4.354 | 1.960 | 2.221 | 0.172 |
| Stomata | Contrast | Estimate | SE | z.ratio | P value |
|  | Microbe-depleted - Rhizobia (MLBS) | 5.362 | 2.499 | 2.146 | 0.201 |
|  | Microbe-depleted - Rhizobia (PP) | *6.624* | *2.658* | *2.492* | *0.092* |
|  | Microbe-depleted - Community (MLBS) | 1.880 | 2.730 | 0.689 | 0.959 |
|  | Microbe-depleted - Community (PP) | 5.017 | 2.985 | 1.681 | 0.446 |
|  | Rhizobia (MLBS) - Rhizobia (PP) | 1.262 | 2.562 | 0.493 | 0.988 |
|  | Rhizobia (MLBS) - Community (MLBS) | -3.483 | 2.637 | -1.321 | 0.678 |
|  | Rhizobia (MLBS) - Community (PP) | -0.345 | 2.900 | -0.119 | 1.000 |
|  | Rhizobia (PP) - Community (MLBS) | -4.744 | 2.788 | -1.702 | 0.433 |
|  | Rhizobia (PP) - Community (PP) | -1.607 | 3.038 | -0.529 | 0.984 |
|  | Community (MLBS) - Community (PP) | 3.137 | 3.102 | 1.012 | 0.850 |
| Trichome density | Contrast | Estimate | SE | z.ratio | P value |
|  | Microbe-depleted - Rhizobia (MLBS) | 10.967 | 2.170 | 5.053 | <0.001 |
|  | Microbe-depleted - Rhizobia (PP) | 9.685 | 2.382 | 4.065 | <0.001 |
|  | Microbe-depleted - Community (MLBS) | -7.429 | 2.380 | -3.122 | 0.016 |
|  | Microbe-depleted - Community (PP) | 9.743 | 2.492 | 3.909 | 0.001 |
|  | Rhizobia (MLBS) - Rhizobia (PP) | -1.282 | 2.176 | -0.589 | 0.977 |
|  | Rhizobia (MLBS) - Community (MLBS) | -18.396 | 2.273 | -8.095 | <0.001 |
|  | Rhizobia (MLBS) - Community (PP) | -1.223 | 2.429 | -0.504 | 0.987 |
|  | Rhizobia (PP) - Community (MLBS) | -17.114 | 2.473 | -6.922 | <0.001 |
|  | Rhizobia (PP) - Community (PP) | 0.059 | 2.686 | 0.022 | 1.000 |
|  | Community (MLBS) - Community (PP) | 17.172 | 2.653 | 6.473 | <0.001 |
| Shoot mass | Contrast | Estimate | SE | z.ratio | P value |
|  | Microbe-depleted - Rhizobia (MLBS) | -0.002 | 0.002 | -1.039 | 0.837 |
|  | Microbe-depleted - Rhizobia (PP) | 0.001 | 0.003 | 0.441 | 0.992 |
|  | Microbe-depleted - Community (MLBS) | 0.013 | 0.002 | 5.526 | <0.001 |
|  | Microbe-depleted - Community (PP) | 0.012 | 0.002 | 5.243 | <0.001 |
|  | Rhizobia (MLBS) - Rhizobia (PP) | 0.004 | 0.002 | 1.717 | 0.423 |
|  | Rhizobia (MLBS) - Community (MLBS) | 0.015 | 0.002 | 6.224 | <0.001 |
|  | Rhizobia (MLBS) - Community (PP) | 0.014 | 0.002 | 5.846 | <0.001 |
|  | Rhizobia (PP) - Community (MLBS) | 0.011 | 0.003 | 4.374 | <0.001 |
|  | Rhizobia (PP) - Community (PP) | 0.011 | 0.003 | 4.169 | <0.001 |
|  | Community (MLBS) - Community (PP) | -0.001 | 0.002 | -0.300 | 0.998 |
| Root mass | Contrast | Estimate | SE | z.ratio | P value |
|  | Microbe-depleted - Rhizobia (MLBS) | -0.002 | 0.001 | -1.152 | 0.779 |
|  | Microbe-depleted - Rhizobia (PP) | 0.002 | 0.002 | 1.277 | 0.706 |
|  | Microbe-depleted - Community (MLBS) | 0.006 | 0.001 | 4.435 | <0.001 |
|  | Microbe-depleted - Community (PP) | 0.007 | 0.001 | 4.936 | <0.001 |
|  | Rhizobia (MLBS) - Rhizobia (PP) | 0.004 | 0.001 | 2.913 | 0.030 |
|  | Rhizobia (MLBS) - Community (MLBS) | 0.008 | 0.001 | 5.245 | <0.001 |
|  | Rhizobia (MLBS) - Community (PP) | 0.008 | 0.001 | 5.609 | <0.001 |
|  | Rhizobia (PP) - Community (MLBS) | *0.004* | *0.002* | *2.624* | *0.066* |
|  | Rhizobia (PP) - Community (PP) | 0.005 | 0.002 | 3.047 | 0.020 |
|  | Community (MLBS) - Community (PP) | 0.001 | 0.002 | 0.372 | 0.996 |
| Shoot:Root | Contrast | Estimate | SE | z.ratio | P value |
|  | Microbe-depleted - Rhizobia (MLBS) | -0.006 | 0.152 | -0.036 | 1.000 |
|  | Microbe-depleted - Rhizobia (PP) | -0.246 | 0.163 | -1.514 | 0.553 |
|  | Microbe-depleted - Community (MLBS) | 0.211 | 0.164 | 1.282 | 0.703 |
|  | Microbe-depleted - Community (PP) | -0.133 | 0.154 | -0.865 | 0.910 |
|  | Rhizobia (MLBS) - Rhizobia (PP) | -0.241 | 0.140 | -1.722 | 0.420 |
|  | Rhizobia (MLBS) - Community (MLBS) | 0.216 | 0.174 | 1.241 | 0.728 |
|  | Rhizobia (MLBS) - Community (PP) | -0.128 | 0.162 | -0.789 | 0.934 |
|  | Rhizobia (PP) - Community (MLBS) | *0.457* | *0.183* | *2.504* | *0.090* |
|  | Rhizobia (PP) - Community (PP) | 0.113 | 0.170 | 0.664 | 0.964 |
|  | Community (MLBS) - Community (PP) | -0.344 | 0.173 | -1.989 | 0.271 |
| Stem Length | Contrast | Estimate | SE | z.ratio | P value |
|  | Microbe-depleted - Rhizobia (MLBS) | 0.219 | 0.178 | 1.233 | 0.732 |
|  | Microbe-depleted - Rhizobia (PP) | 0.468 | 0.193 | 2.425 | 0.109 |
|  | Microbe-depleted - Community (MLBS) | 1.227 | 0.168 | 7.284 | <0.001 |
|  | Microbe-depleted - Community (PP) | 1.239 | 0.169 | 7.314 | <0.001 |
|  | Rhizobia (MLBS) - Rhizobia (PP) | 0.248 | 0.152 | 1.631 | 0.477 |
|  | Rhizobia (MLBS) - Community (MLBS) | 1.008 | 0.179 | 5.620 | <0.001 |
|  | Rhizobia (MLBS) - Community (PP) | 1.019 | 0.187 | 5.463 | <0.001 |
|  | Rhizobia (PP) - Community (MLBS) | 0.760 | 0.198 | 3.837 | <0.001 |
|  | Rhizobia (PP) - Community (PP) | 0.771 | 0.197 | 3.915 | <0.001 |
|  | Community (MLBS) - Community (PP) | 0.011 | 0.188 | 0.060 | 1.000 |
| Number of nodules | Contrast | Estimate | SE | z.ratio | P value |
|  | Microbe-depleted - Rhizobia (MLBS) | -11.453 | 1.935 | -5.920 | <0.001 |
|  | Microbe-depleted - Rhizobia (PP) | -12.036 | 2.155 | -5.586 | <0.001 |
|  | Microbe-depleted - Community (MLBS) | -7.880 | 2.013 | -3.915 | 0.001 |
|  | Microbe-depleted - Community (PP) | -9.376 | 2.055 | -4.563 | <0.001 |
|  | Rhizobia (MLBS) - Rhizobia (PP) | -0.583 | 1.906 | -0.306 | 0.998 |
|  | Rhizobia (MLBS) - Community (MLBS) | 3.573 | 1.963 | 1.821 | 0.361 |
|  | Rhizobia (MLBS) - Community (PP) | 2.077 | 2.035 | 1.021 | 0.846 |
|  | Rhizobia (PP) - Community (MLBS) | 4.156 | 2.185 | 1.902 | 0.316 |
|  | Rhizobia (PP) - Community (PP) | 2.660 | 2.206 | 1.206 | 0.748 |
|  | Community (MLBS) - Community (PP) | -1.496 | 2.155 | -0.694 | 0.958 |

**Table S5: ANOVA results for the growth and nodulation relationships tested in Figure 4.** We tested the significance of growth PC1 and nodulation traits in explaining variation in leaf area and trichome density across microbial treatments.

|  | **Chi-squared** | **Degrees of freedom** | **P-value** |
| --- | --- | --- | --- |
| **Leaflet area** | | | |
| **Growth and leaflet area** | | | |
| Microbial treatment | 5.395 | 4 | 0.2491 |
| Growth PC1 | 38.578 | 1 | <0.001 |
| Microbial treatment:Growth PC1 | 6.829 | 4 | 0.1452 |
| **Nodulation and leaflet area** | | | |
| Microbial treatment | 103.640 | 3 | <0.001 |
| Number of nodules (z-scored) | 6.241 | 1 | 0.0125 |
| Microbial treatment:Nodules (z-scored) | 2.789 | 3 | 0.4253 |
| **Trichome density** | | | |
| **Growth and trichome density** | | | |
| Microbial treatment | 15.903 | 4 | 0.0032 |
| Growth PC1 | 1.753 | 1 | 0.1855 |
| Microbial treatment:Growth PC1 | 21.999 | 4 | <0.001 |
| **Nodulation and trichome density** | | | |
| Microbial treatment | 51.403 | 3 | <0.001 |
| Number of nodules (z-scored) | 2.924 | 1 | 0.0873 |
| Microbial treatment:Nodules (z-scored) | 8.337 | 3 | 0.0395 |

**Table S6: Post-hoc tests comparing treatment-driven differences in the relationship between trichome density and growth or nodulation.**

| **Trichomes and growth** | | | | | |
| --- | --- | --- | --- | --- | --- |
| Contrast | estimate | SE | df | z.ratio | p.value |
| Microbe-depleted - Rhizobia (MLBS) | 3 | 1.515 | Inf | 2.292 | 0.147 |
| Microbe-depleted - Rhizobia (PP) | 0 | 1.660 | Inf | 0.189 | 1.000 |
| Microbe-depleted - Community (MLBS) | **13** | **3.184** | **Inf** | **3.981** | **0.001** |
| Microbe-depleted - Community (PP) | -3 | 3.855 | Inf | -0.673 | 0.962 |
| Rhizobia (MLBS) - Rhizobia (PP) | -3 | 1.492 | Inf | -2.116 | 0.213 |
| Rhizobia (MLBS) - Community (MLBS) | **9** | **3.116** | **Inf** | **2.954** | **0.026** |
| Rhizobia (MLBS) - Community (PP) | -6 | 3.805 | Inf | -1.595 | 0.501 |
| Rhizobia (PP) - Community (MLBS) | **12** | **3.188** | **Inf** | **3.877** | **0.001** |
| Rhizobia (PP) - Community (PP) | -3 | 3.864 | Inf | -0.753 | 0.944 |
| Community (MLBS) - Community (PP) | **-15** | **4.726** | **Inf** | **-3.231** | **0.011** |
| **Trichomes and nodules** | | | | | |
| Contrast | estimate | SE | df | z.ratio | p.value |
| Rhizobia (MLBS) - Rhizobia (PP) | -0.410 | 0.236 | Inf | -1.734 | 0.306 |
| Rhizobia (MLBS) - Community (MLBS) | 0.689 | 0.419 | Inf | 1.647 | 0.352 |
| Rhizobia (MLBS) - Community (PP) | -0.542 | 0.413 | Inf | -1.313 | 0.554 |
| Rhizobia (PP) - Community (MLBS) | **1.099** | **0.442** | **Inf** | **2.489** | **0.062** |
| Rhizobia (PP) - Community (PP) | -0.132 | 0.436 | Inf | -0.303 | 0.990 |
| Community (MLBS) - Community (PP) | -1.232 | 0.557 | Inf | -2.213 | 0.120 |

**Figure S3: The relationship between nodule number and growth, or the benefit per nodule overall and for each microbial treatment.** ANOVA results in Table S7.

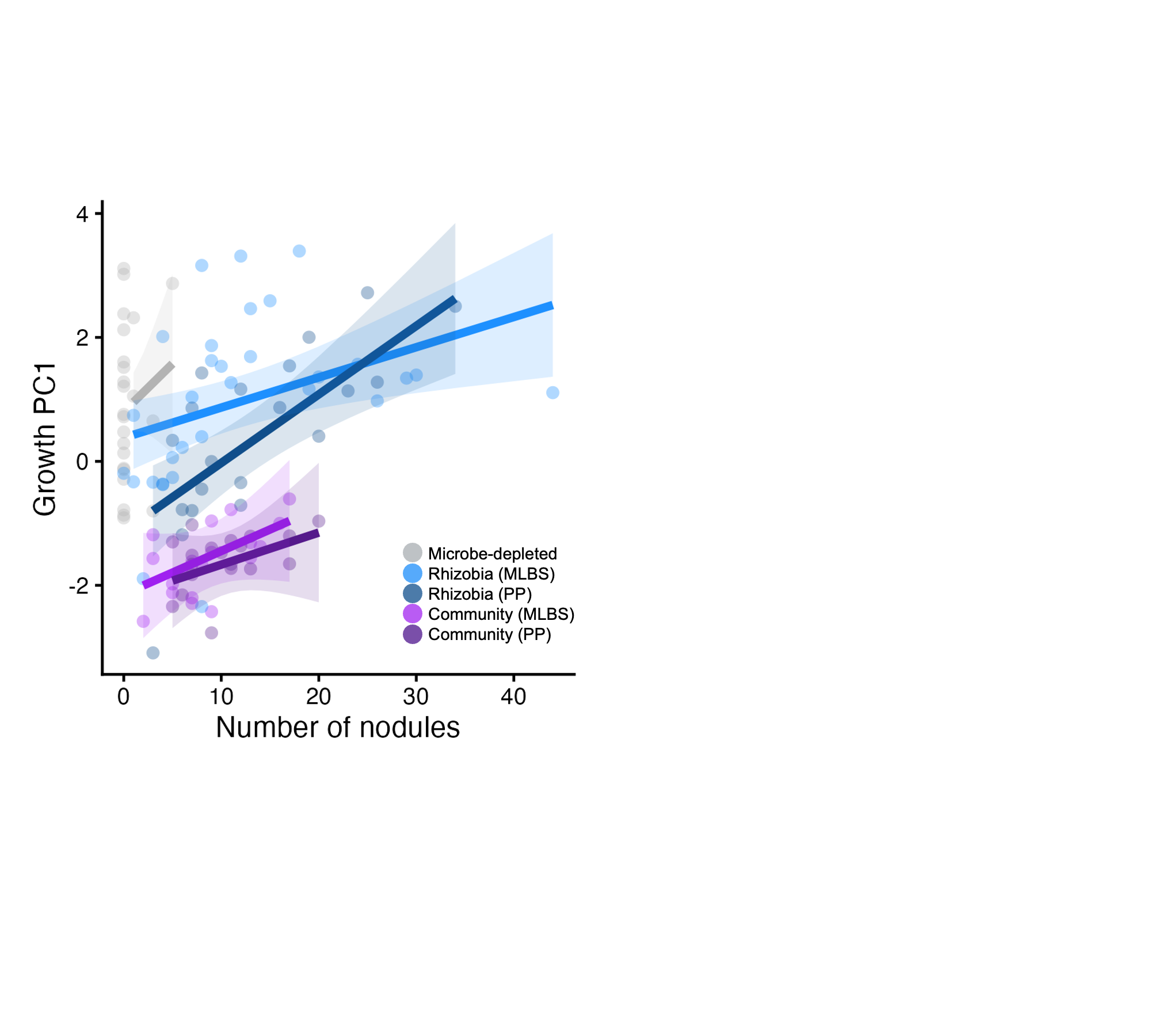

**Table S7: ANOVA results for the relationship of nodulation, microbial treatment, and the interaction of microbial treatment and nodule number on overall growth.**

|  | **Chisq** | **Df** | **P-value** |
| --- | --- | --- | --- |
| **Microbial treatment** | 77.6595 | 4 | **<0.001** |
| **Nodules (z-scored)** | 6.0384 | 1 | **0.0140** |
| **Treatment:Nodules** | 4.1671 | 4 | 0.3839 |

**Figure S4: Subsetted principal components and linear discriminant analyses for leaf traits excluding trichome density.** (A-E) Principal component analyses of the four leaf traits excluding trichome density. (F-H) Comparisons of microbial treatments against the microbe-depleted control using **P-**matrices composed of the same four leaf traits. (I) Linear discriminant analysis for the four leaf traits. Tests robustness of results to exclusion of trichome density (Tables S8-11).
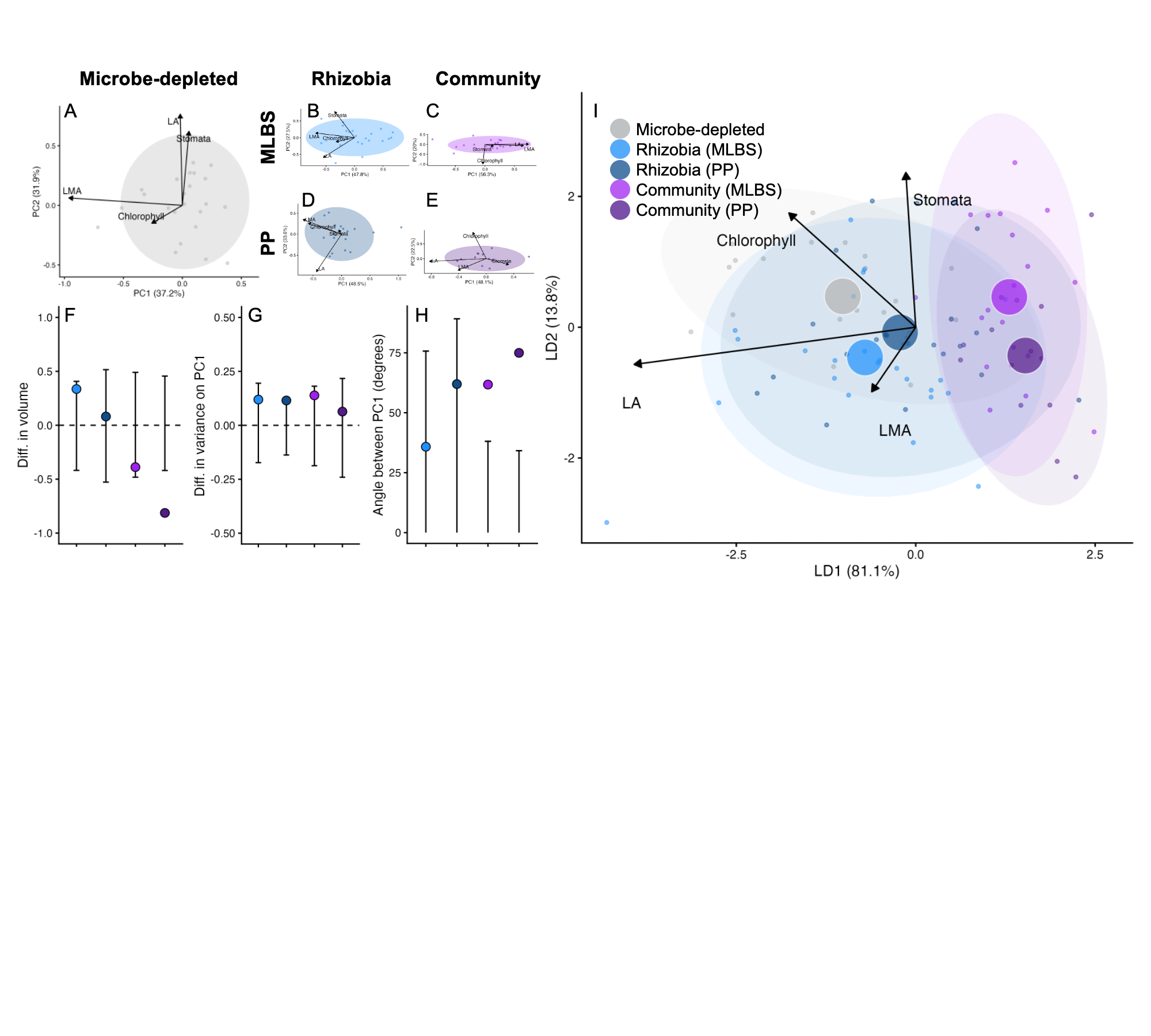

**Table S8: MANOVA results for all trait suites (all leaf traits, growth traits, and leaf traits excluding trichome density), testing overall multivariate treatment effects.**

|  | **Df** | **Pillai** | **approx F** | **num Df** | **den Df** | **P-value** |
| --- | --- | --- | --- | --- | --- | --- |
| **Leaf traits** | 4 | 1.120 | 7.468 | 20 | 384 | <0.001 |
| **Growth traits** | 4 | 0.764 | 6.377 | 16 | 432 | <0.001 |
| **Leaf traits without trichome density** | 4 | 0.729 | 5.407 | 16 | 388 | <0.001 |

**Table S9: P-matrix comparisons and permutation null distribution ranges for leaf traits excluding trichome density**. Companion to Figure S3 panels F-H: provides values and significance thresholds for matrix comparisons for leaf traits when trichome density is excluded.

|  | **Rhizobia (MLBS)** | **Rhizobia (PP)** | **Community (MLBS)** | **Community (PP)** |
| --- | --- | --- | --- | --- |
| Volume | 0.4115 | 0.2050 | 0.1265 | -0.2777 |
| *Volume (95% CI)* | *-0.464 — 0.4457* | *-0.5781 — 0.5687* | *-0.5323 — 0.5362* | *-0.4504 — 0.4779* |
| Proportion of Pmax | 0.1058 | 0.1131 | **0.1906** | 0.1086 |
| *Proportion of Pmax (95% CI)* | *-0.1778 — 0.1997* | *-0.1335 — 0.106* | *-0.1802 — 0.1751* | *-0.2329 — 0.2221* |
| Angle between Pmax | 42.74 | 35.74 | **44.97** | **61.21** |
| *Angle between Pmax (95% CI)* | *75.23* | *89.14* | *36.02* | *33.12* |

**Table S10: Trait loadings for treatment-specific principal components analyses (on covariance matrices) for leaf traits excluding trichome density.**

|  | **Microbe-depleted** | **Rhizobia (MLBS)** | **Rhizobia (PP)** | **Community (MLBS)** | **Community (PP)** |
| --- | --- | --- | --- | --- | --- |
| **PropVar** | 0.3723 | 0.4781 | 0.4854 | 0.5629 | 0.4810 |
| Leaflet Area | -0.0183 | -0.5512 | -0.4427 | 0.6551 | -0.8273 |
| Leaflet Mass Area | -0.9638 | -0.6840 | -0.6790 | 0.7400 | -0.4121 |
| Chlorophyll | -0.2600 | -0.3257 | -0.5837 | -0.0368 | -0.1960 |
| Stomata density | 0.0561 | -0.3495 | -0.0476 | 0.1478 | 0.3276 |

**Table S11: Trait loadings for leaf traits excluding trichome density on linear discriminant analysis and principal components analyses, with the proportion of trace and variance explained by each axis.** Companion to Figure S3 panel I.

| **Linear Discriminant Analysis** | | | | |
| --- | --- | --- | --- | --- |
|  | **LD1** | **LD2** | **LD3** | **LD4** |
| *Proportion of Trace* | *0.811* | *0.138* | *0.048* | *0.002* |
| Leaflet Area | -0.981 | -0.142 | 0.091 | 0.098 |
| Leaflet Mass Area | -0.153 | -0.247 | -0.246 | 0.925 |
| Chlorophyll | -0.442 | 0.437 | -0.741 | 0.255 |
| Stomata Density | -0.034 | 0.592 | 0.645 | 0.482 |
| **Global Principal Components Analysis (across all treatments)** | | | | |
|  | **PC1** | **PC2** | **PC3** | **PC4** |
| *Proportion of Variance* | *0.4124* | *0.2637* | *0.1872* | *0.1368* |
| Leaf Area | -0.5395 | -0.0382 | -0.8068 | -0.2377 |
| Leaf Mass Area | -0.5825 | 0.1894 | 0.5483 | -0.5693 |
| Chlorophyll | -0.5992 | -0.3092 | 0.2065 | 0.7090 |
| Stomata Density | -0.1026 | 0.9312 | -0.0761 | 0.3415 |

**Table S12: Correlation (bottom left) and covariance (top right) matrices for leaf (A) and growth (B) traits across all treatments.** Diagonals show trait variances.

|  | **A) Leaf traits** | | | | | |  | **B) Growth traits** | | | | | |
| --- | --- | --- | --- | --- | --- | --- | --- | --- | --- | --- | --- | --- | --- |
|  | **Microbe-depleted** | | | | | | | | | | | | |
|  |  | | *cov* | | | |  |  | | | *cov* | | |
|  | *variance* | **Leaflet Area** | **LMA** | **Chlorophyll** | **Stomata** | **Trichomes** |  |  | *variance* | **Shoot (g)** | **Root (g)** | **Shoot:Root** | **Longest Stem** |
|  | **Leaflet Area** | 0.012 | <0.001 | 0.145 | 0.257 | 0.184 |  |  | **Shoot (g)** | <0.001 | <0.001 | <0.001 | 0.002 |
| *cor* | **LMA** | -0.025 | <0.001 | 0.001 | 0.001 | <0.001 |  | *cor* | **Root (g)** | 0.881 | <0.001 | -0.001 | 0.001 |
|  | **Chlorophyll** | 0.244 | 0.238 | 30.859 | -33.113 | 0.021 |  |  | **Shoot:Root** | 0.069 | -0.348 | 0.201 | -0.019 |
|  | **Stomata** | 0.247 | 0.068 | -0.613 | 94.699 | 9.986 |  |  | **Longest Stem** | 0.421 | 0.454 | -0.089 | 0.237 |
|  | **Trichomes** | 0.253 | 0.055 | 0.001 | 0.151 | 45.953 |  |  |  |  |  |  |  |
|  | **Rhizobia (MLBS)** | | | | | | | | | | | | |
|  |  | | *cov* | | | |  |  | | | *cov* | | |
|  | *variance* | **Leaflet Area** | **LMA** | **Chlorophyll** | **Stomata** | **Trichomes** |  |  | *variance* | **Shoot (g)** | **Root (g)** | **Shoot:Root** | **Longest Stem** |
|  | **Leaflet Area** | 0.018 | <0.001 | 0.206 | 0.052 | -0.096 |  |  | **Shoot (g)** | <0.001 | <0.001 | 0.001 | 0.003 |
| *cor* | **LMA** | 0.384 | <0.001 | 0.003 | 0.004 | -0.005 |  | *cor* | **Root (g)** | 0.872 | <0.001 | -0.001 | 0.002 |
|  | **Chlorophyll** | 0.245 | 0.545 | 40.141 | 0.191 | -15.441 |  |  | **Shoot:Root** | 0.169 | -0.296 | 0.171 | 0.004 |
|  | **Stomata** | 0.033 | 0.372 | 0.003 | 138.025 | -4.230 |  |  | **Longest Stem** | 0.683 | 0.646 | 0.021 | 0.229 |
|  | **Trichomes** | -0.096 | -0.617 | -0.323 | -0.048 | 56.962 |  |  |  |  |  |  |  |
|  | **Rhizobia (PP)** | | | | | | | | | | | | |
|  |  | | *cov* | | | |  |  | | | *cov* | | |
|  | *variance* | **Leaflet Area** | **LMA** | **Chlorophyll** | **Stomata** | **Trichomes** |  |  | *variance* | **Shoot (g)** | **Root (g)** | **Shoot:Root** | **Longest Stem** |
|  | **Leaflet Area** | 0.013 | <0.001 | 0.139 | -0.046 | 0.307 |  |  | **Shoot (g)** | <0.001 | <0.001 | <0.001 | 0.003 |
| *cor* | **LMA** | 0.125 | <0.001 | 0.005 | 0.000 | -0.002 |  | *cor* | **Root (g)** | 0.833 | <0.001 | -0.001 | 0.002 |
|  | **Chlorophyll** | 0.162 | 0.758 | 58.162 | 7.847 | -9.591 |  |  | **Shoot:Root** | 0.022 | -0.478 | 0.379 | -0.093 |
|  | **Stomata** | -0.059 | 0.067 | 0.149 | 47.855 | -0.421 |  |  | **Longest Stem** | 0.615 | 0.657 | -0.232 | 0.425 |
|  | **Trichomes** | 0.375 | -0.302 | -0.173 | -0.008 | 52.674 |  |  |  |  |  |  |  |
|  | **Community (MLBS)** | | | | | | | | | | | | |
|  |  | | *cov* | | | |  |  | | | *cov* | | |
|  | *variance* | **Leaflet Area** | **LMA** | **Chlorophyll** | **Stomata** | **Trichomes** |  |  | *variance* | **Shoot (g)** | **Root (g)** | **Shoot:Root** | **Longest Stem** |
|  | **Leaflet Area** | 0.004 | <0.001 | -0.010 | 0.076 | -0.463 |  |  | **Shoot (g)** | <0.001 | <0.001 | <0.001 | <0.001 |
| *cor* | **LMA** | 0.580 | <0.001 | <0.001 | 0.002 | -0.007 |  | *cor* | **Root (g)** | 0.852 | <0.001 | -0.001 | <0.001 |
|  | **Chlorophyll** | -0.026 | -0.050 | 36.608 | 2.086 | -17.386 |  |  | **Shoot:Root** | -0.142 | -0.604 | 0.182 | -0.038 |
|  | **Stomata** | 0.182 | 0.322 | 0.050 | 47.117 | -26.810 |  |  | **Longest Stem** | 0.274 | 0.264 | -0.258 | 0.119 |
|  | **Trichomes** | -0.686 | -0.689 | -0.260 | -0.354 | 121.760 |  |  |  |  |  |  |  |
|  | **Community (PP)** | | | | | | | | | | | | |
|  |  | | *cov* | | | |  |  | | | *cov* | | |
|  | *variance* | **Leaflet Area** | **LMA** | **Chlorophyll** | **Stomata** | **Trichomes** |  |  | *variance* | **Shoot (g)** | **Root (g)** | **Shoot:Root** | **Longest Stem** |
|  | **Leaflet Area** | 0.003 | <0.001 | 0.048 | -0.084 | 0.093 |  |  | **Shoot (g)** | <0.001 | <0.001 | 0.001 | 0.001 |
| *cor* | **LMA** | 0.434 | <0.001 | <0.001 | -0.002 | <0.001 |  | *cor* | **Root (g)** | 0.448 | <0.001 | <0.001 | <0.001 |
|  | **Chlorophyll** | 0.196 | -0.097 | 20.354 | -6.954 | -2.313 |  |  | **Shoot:Root** | 0.478 | -0.522 | 0.400 | 0.126 |
|  | **Stomata** | -0.209 | -0.376 | -0.207 | 55.716 | -0.918 |  |  | **Longest Stem** | 0.763 | 0.175 | 0.547 | 0.133 |
|  | **Trichomes** | 0.258 | -0.110 | -0.077 | -0.018 | 44.264 |  |  |  |  |  |  |  |
